# Distinct prelimbic cortex neuronal responses to emotional states of others drive emotion recognition in adult male mice

**DOI:** 10.1101/2024.03.03.583184

**Authors:** Renad Jabarin, Alok Nath Mohapatra, Natali Ray, Shai Netser, Shlomo Wagner

## Abstract

The ability to perceive the emotional states of others, termed emotion recognition, allows individuals to adapt their conduct to the social environment. The brain mechanisms underlying this capacity, known to be impaired in individuals with autism spectrum disorder (ASD), remain, however, elusive. Using the emotional state preference paradigm, we show that adult mice can discern emotional states of conspecifics in a sex-specific manner, behavior impaired in an ASD mouse model. Fiber photometry revealed inhibition of pyramidal neurons in the prelimbic medial prefrontal cortex (PrL) during investigation of aroused individuals, as opposed to transient excitation towards non-aroused conspecifics. Augmenting this differential neuronal response enhanced emotion recognition, while abolishing this response eliminated such behavior, as demonstrated via optogenetic stimulation. Chronic electrophysiological recordings at the single-cell level indicated social stimulus-specific responses in PrL neurons at the onset and conclusion of social investigation bouts, potentially regulating the initiation and termination of social interactions. The dysregulated activity of these neurons may thus contribute to deficits in social behavior observed in ASD.

## Background

Social cognition involves the perceiving and interpreting of social cues exchanged between individuals, which are imperative for the adaptive response of a subject to the social milieu ^1,2^. Emotion recognition, corresponding to the ability to discern the emotional states of others ^3–5^, plays a pivotal role in various pro-social behaviors, such as emotion contagion, empathy, and helping behavior ^6–8^. This capacity, which allows an individual to adapt its behavior to the emotional states of others, is known to be compromised in individuals diagnosed with autism spectrum disorder (ASD) ^9–11^. Despite the extensive insight gained from functional magnetic resonance imaging (fMRI) studies that identified brain areas active during emotion recognition tasks ^4,12–15^, the specific deficits in brain activity and neuronal network dynamics underlying impaired emotion recognition in ASD remain elusive.

Animal models of ASD offer a valuable experimental system with which to investigate such deficits, allowing for invasive monitoring and manipulation of neural activity during social behavior ^16,17^. However, until recently, reliable behavioral tasks for assessing emotion recognition in animal models were lacking. Recent studies, including our own, have demonstrated that mice possess the ability to discriminate between conspecifics based on their emotional states ^18–20^, thus providing the means to evaluate emotion recognition in murine models of ASD and other neurodevelopmental disorders ^21^. This observation led us and others to develop a novel behavioral paradigm, which we named “emotional-state preference” (ESP). In this task, a subject mouse is presented with two conspecifics (stimulus animals), one of which has undergone emotional arousal manipulation. Using one version of this task, we previously demonstrated that mice show a preference for investigating socially isolated conspecifics over group-housed stimulus animals ^19^. In the same study, we also demonstrated the specific deficiency of mice expressing the A350V-encoding mutation in the *Iqsec2* gene, a mutation associated with ASD in humans, in this task ^22^. Relying on two other versions of this task, Ferretti *et al*. reported that C57BL/6J mice exhibited a preference for investigating stressed or relieved conspecifics over their neutral counterparts ^18^. Here, we performed these three versions of the ESP paradigm in both male and female C57BL/6J mice and showed that male mice could discriminate between distinct states of arousal. Moreover, we found that male *Shank3*-deficient mice, a well-established murine ASD model, exhibit impaired ESP behavior. These observations further suggest that the ESP behavioral paradigm is a valid model for emotion recognition.

To date, only few studies have explored the brain mechanisms underlying emotion recognition in mice. Such studies pointed to oxytocin signaling in the central amygdala and GABAergic inhibitory interneurons in the medial prefrontal cortex (mPFC) as neuronal mechanisms involved in the recognition of stress and relieved states by mice ^18,20^. Notably, the mPFC was previously implicated in social decision making in both humans and animal models ^23,24^. Here, we recorded extracellular electrophysiological signals from multiple social behavior-associated brain regions, and identified the prelimbic (PrL) region of the mPFC as exhibiting differential rhythmic population activity during investigations of stressed vs. neutral stimulus animals, thus suggesting involvement of this region in ESP behavior. Using fiber photometry, we confirmed that PrL pyramidal neurons exhibit differential responses during investigation of aroused vs. non-aroused (neutral) stimulus animals. By optogenetically exciting these cells, we found that brief excitation at the beginning of stimulus investigation affected the subject’s behavior during ESP in a stimulus animal-specific manner. Finally, we made use of chronically implanted Neuropixels 1.0 probes to monitor the activity of hundreds of mPFC neurons in behaving mice and identified multiple groups of neurons which responded specifically at the onset or end of investigation bouts towards specific stimuli. Importantly, some of these neuron groups started to change their activity about one second before the behavioral event, suggesting that PrL pyramidal neurons participate in the decision of whether to initiate or terminate investigation of a given stimulus. Together, our results suggest that differential responses of PrL pyramidal neurons to aroused vs. neutral conspecifics is crucial for emotion recognition and that these neurons participate in socio-emotional decision-making processes.

## Results

### Testing male and female mice with three versions of the ESP paradigm

To test mice with several different versions of the ESP paradigm, we first followed the study of Ferretti *et al.* ^18^ to examine and validate the behavior of adult C57BL/6J mice in the relief version of the ESP task (**ESPr, Fig. 1A**). As anticipated, we found that both male and female mice investigated the relieved stimulus animal for significantly more time than spent with a neutral mouse, thus exhibiting a relief-state preference (**Fig. 1B-C**). In agreement with our previous results using other social discrimination tasks ^19,25,26^, the preference for a specific stimulus animal by C57BL/6J subjects was reflected only during long (>6 s), but not short (≤6 s) investigation bouts (**Fig. S1A, B**). We observed similar results with the stress version of ESP (**ESPs**), where both male and female subjects preferred investigating a stressed stimulus animal over a neutral peer (**Fig. 1D-F, Fig. S1C-D**). Finally, using our previously established isolation version of ESP (**ESPi, Fig. 1G**) ^19^, we found that male mice preferred to investigate an isolated mouse over a group-housed animal (**Fig. 1H, Fig. S1E**). Females, however, did not show such preference in this task (**Fig. 1I, Fig. S1F**), even when male mice were used as stimulus animals (**Fig. S1G-H**). Thus, unlike the mouse behavior seen in the ESPr and ESPs tasks, performance in the ESPi task seems to be sex-specific, being exhibited only by male mice.

Our findings showing sex-specific behavior solely in the ESPi task suggests that stimulus mice in a state of isolation differ from mice in the relief and stress states. We, therefore, examined the preferences of male and female subjects while simultaneously exposing them to three stimulus animals, each in a distinct arousal state (i.e., isolation, relief, or stress) and each located in a different chamber within the same arena (**Fig. 1J**). We found that both male and female mice spent significantly less time investigating the isolated stimulus, as compared to the two other stimuli (**Fig. 1K-L, Fig. S1I-J**). While this result was expected for females, which did not prefer isolated over group-housed stimulus animals (Fig. 1F), our finding that males also showed differential preference for the various stimulus animals once again suggests that they can discriminate between distinct emotional states. These results thus support the validity of the ESP paradigm as a model for studying emotion recognition.

**Figure 1.**
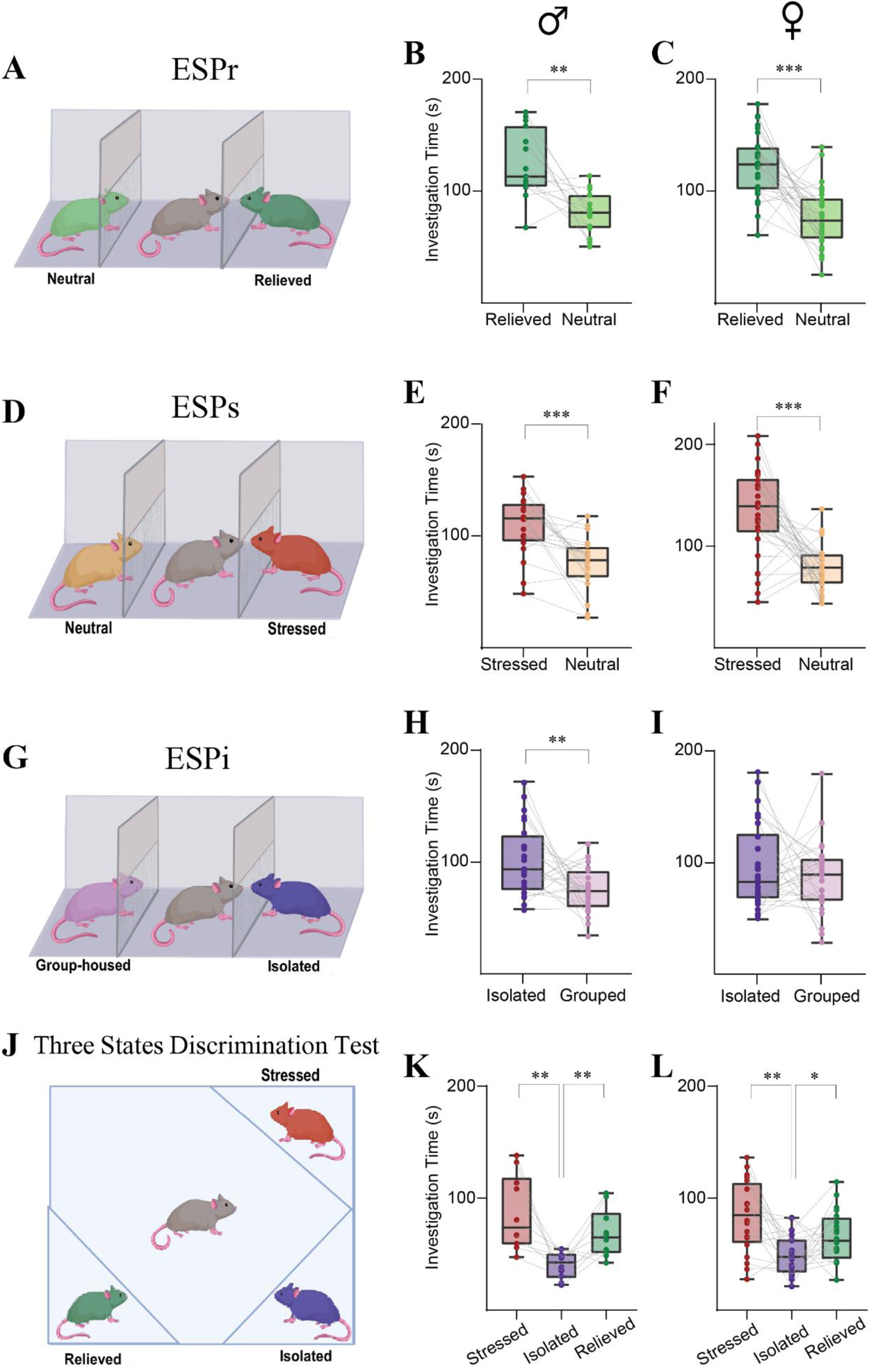
Mice can distinguish the emotional states of others. **A.** Schematic representation of the ESPr task. **B.** Mean time dedicated by male mice for investigating relieved (green) or neutral (light green) stimulus animals during the ESPr task (n=15 subjects). **C.** As in **B**, for female mice (n=30 subjects). **D-F**. As in **A-C**, for the ESPs task (n_males_=20 subjects, n_females_=30 subjects). **G-I**. As in **A-C**, for the ESPi task (n_males_=24 subjects, n_females_=30 subjects). **J**. Schematic representation of the arena containing all three types of aroused stimulus animals, as noted above or below the chambers. **K.** Mean time dedicated by male mice (n=10 subjects) for investigating each of the three stimulus animals shown in **J**. **L.** As in **K**, for female subjects (n=20 subjects). B-I: \*\**p*<0.01, \*\*\**p*<0.001, paired samples t-test; L-K: \*\**p*<0.01, \**p*<0.05, *post-hoc* paired LSD comparisons following main effect (*p*<0.01) in one-way repeated measures (RM) ANOVA. The box plot represents 25 to 75 percentiles of the distribution, while the bold line is the median of the distribution. Whiskers represent the smallest and largest values in the distribution.

### *Shank3*-knockout (KO) mice do not exhibit emotional state preference

Since human individuals diagnosed with ASD are known to exhibit impaired emotion recognition^9–11^, we examined whether *Shank3*-KO mice, a well-established genetic mouse model for ASD ^27^, are also impaired in their emotion recognition ability. To that end, we used the ESPs task, which we found to be the most robust and consistent version of ESP (**Fig. 1K-L**). We also tested these mice in three other social discrimination tasks we previously described ^19^, which do not rely on the emotional state of the stimulus animal. These included Social Preference (SP), Sex Preference (SxP) and Social Novelty Preference (SNP) tasks. We found that both wild-type (WT) and Shank3^-/-^ (KO) male mice exhibited a clear preference in the SP and SxP tasks (**Fig. 2A-F**). In contrast, only WT mice exhibited a preference in the SNP and ESPs tasks (**Fig. 2G-L**). No difference between the two genotypes was found in the distance traveled during any of the tasks (**Fig. 2C, F, I, L**), indicative of no change in their motor activity. These findings, together with our previous studies using two other murine models of ASD (i.e., *Iqsec2* A350V ^19^ and *Cntnap2*-KO mice ^28^), suggest that ESP may be especially sensitive to ASD-associated mutations, as this was the only task in which all ASD models showed impairment. Overall, these results further suggest that the ESP paradigm can serve as a valid animal model for human emotion recognition.

**Figure 2.**
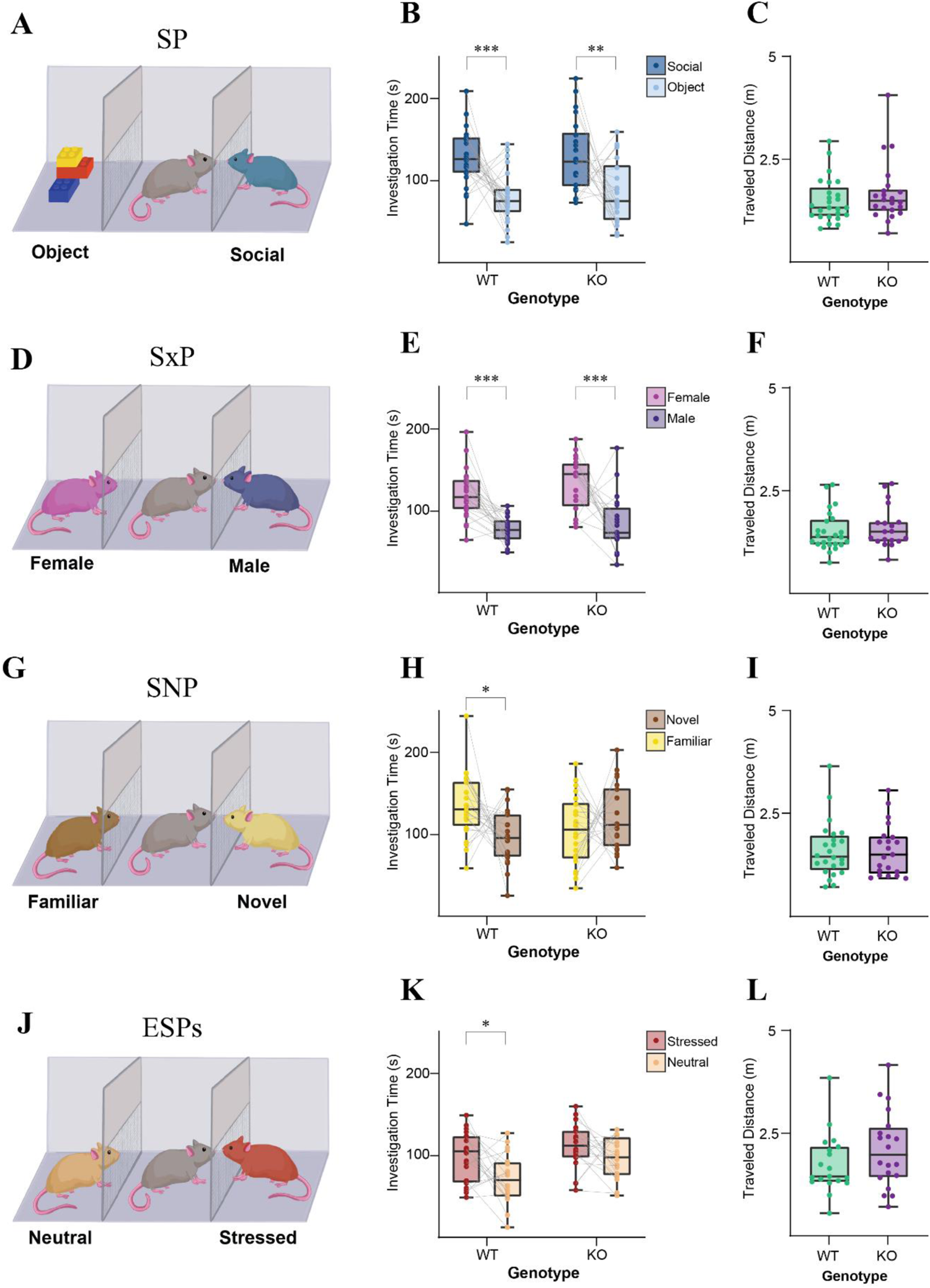
Shank3-KO mice are impaired in emotion recognition. **A.** Schematic representation of the Social Preference (SP) task. **B.** Mean time dedicated by wild-type (WT; left, n=25 subjects) and *Shank3*-KO (KO; right, n=22 subjects) male mice for investigating the social (dark blue) or object (light blue) stimulus during the SP task. A significant main effect was found in two-way mixed-model (MM) ANOVA for stimulus (*p*<0.001). **C.** Mean distance travelled by WT (left) and KO (right) subjects during the SP task. **D-F.** As in **A-C**, for the SxP task (n_WT_=26 subjects, n_KO_= 20 subjects). Significant main effects were revealed by a two-way MM ANOVA for stimulus (*p*<0.001) and genotype (*p*<0.05). **G-I.** As in **A-C**, for the SNP task (n_WT_=24 subjects, n_KO_= 22 subjects). A significant main effect was revealed by a two-way MM ANOVA for stimulus X genotype interaction (*p*<0.05). **J-L.** As in **A-C**, for the ESP task (n_WT_=20 subjects, n_KO_= 20 subjects). Significant main effects were revealed by a two-way MM ANOVA for stimulus (*p*<0.01) and genotype (*p*<0.01). \**p*<0.05, \*\**p*<0.01, \*\*\**p*<0.001, *post-hoc* paired t-test with Holm-Šídák correction for multiple comparisons following the identification of main effects by ANOVA.

### Local field potential rhythmicity links ESP behavior to the medial prefrontal cortex

To screen for brain regions involved in ESP, we used chronically-implanted electrode arrays (**Fig. 3A-B**) ^29^ to record local field potential (LFP) signals from eight social behavior-associated forebrain areas (**Fig. 3C**) in male mice conducting the ESPi task (**Fig. 3D**). We first calculated the change (Δ) in theta and gamma power of the LFP signals recorded during the five min-long encounter, as compared to the five min baseline (pre-encounter) period before introducing stimuli into the arena. We then used Z-score analysis (across the five seconds before and after the start of each investigation bout) to examine changes in theta (**Fig. 3E**) and gamma (**Fig. 3F**) powers, specifically at the beginning of long investigation bouts towards the isolated or group-housed stimulus animals. Of the eight recorded areas, only the PrL region of the mPFC showed a statistically significant difference (corrected for multiple comparisons following a main effect revealed by ANOVA) between the isolated and group-housed stimulus animals, which was specific to gamma power (**Fig. 3E-F**). Interestingly, gamma power was higher during investigation of group-housed, as compared to isolated stimulus animals, and this difference was observed mainly during the first three seconds of the investigation session (**Fig. 3G-I**). Overall, these results point to an association between the PrL and emotion recognition ability in mice.

**Figure 3.**
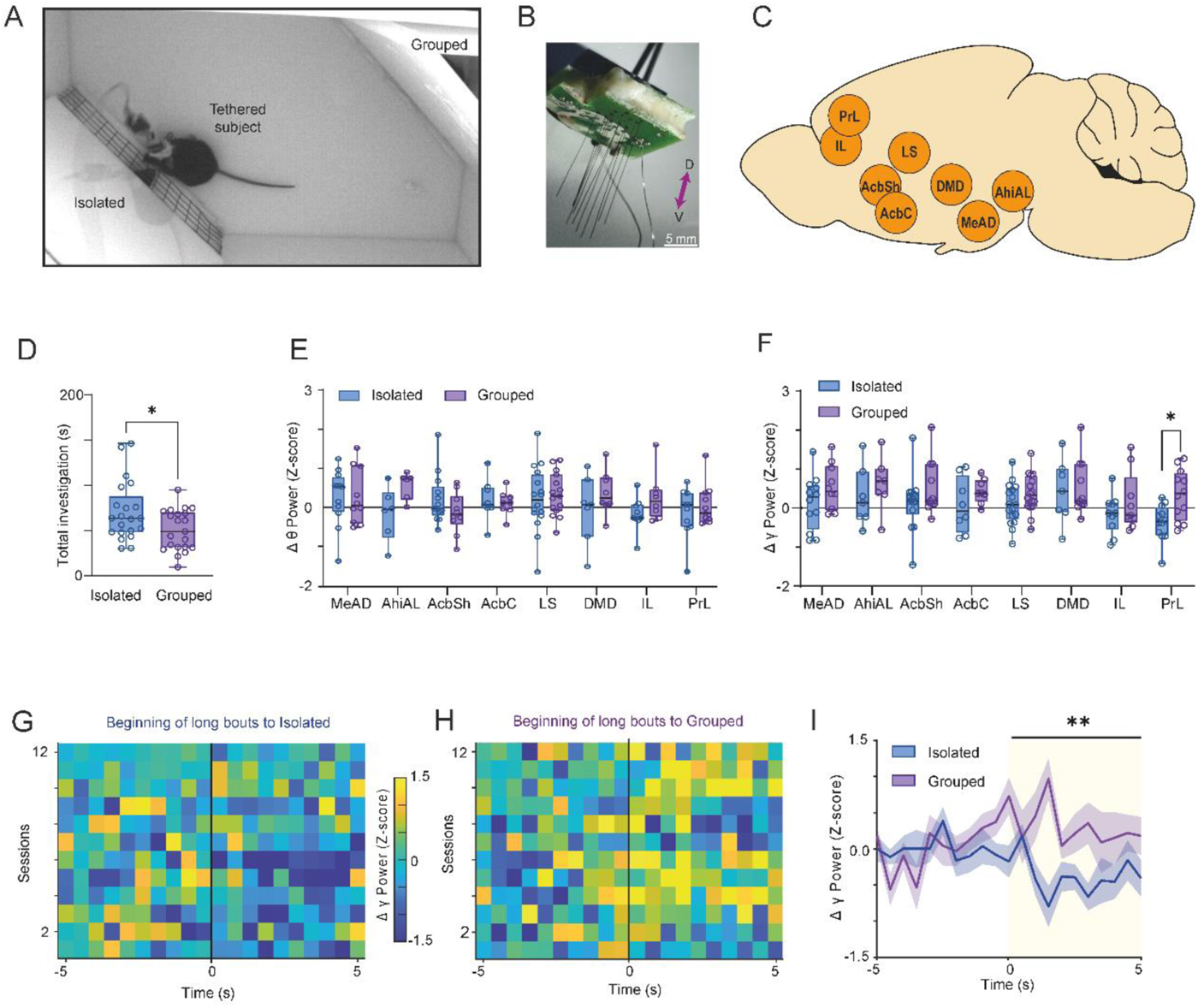
Population activity in the social brain reveals specific involvement of the PrL in emotional state preference. **A.** A picture taken from above of a recorded subject mouse in the arena during the ESPi task. **B.** A picture of a microelectrode array (EAr). **C.** Schematic representation of the eight recorded brain areas analyzed: Medial prefrontal cortex prelimbic (PrL, n=12_isolated_/12_grouped_) and infralimbic (IL, n=10/9) areas, nucleus accumbens shell (AcbSh, n=14/11) and core (AcbC, n=8/8), lateral septum (LS, n=17/16), dorsomedial hypothalamic nucleus (DMD, n=7/11), medial amygdala (MeAD, n=12/10) and amygdalo-hippocampal area, anterolateral part (AhiAL, n=7/7). **D.** Median time dedicated by recorded male subjects for investigating the isolated (blue) or neutral (purple) stimulus animals during the ESPi task (n=15 subjects). \**p* = 0.0351, Wilcoxon test. **E.** Median change (Δ) in theta power, Z-scored across the first five seconds of long (>6 s) bouts using the five seconds before the bout as a baseline, for the eight brain regions simultaneously recorded during ESPi sessions. None of the brain regions showed a statistically significant difference between investigation bouts towards the grouped (purple) or isolated (blue) stimulus animals. Mixed-effects model (REML). Regions x stimulus, regions: F (7, 155) = 0.3561, *p* = 0.9261; stimulus: F (1, 155) = 6.726, *p* = 0.0104; interaction: F (7, 155) = 0.7195, *p* = 0.6556. **F.** As in **E**, for gamma power. Note that the PrL showed a significant difference between investigation bouts towards the grouped and isolated stimulus animals. Mixed-effects model (REML). Regions x stimulus, regions: F (7, 155) = 2.111, *p* = 0.1282; stimulus: F (1, 155) = 13, *p* = 0.0004; interaction: F (7, 155) = 0.4209, *p* = 0.8882, Šídák’s *post-hoc* test, PrL: t (155) =2.774, \**p* = 0.0487. **G.** Heat-map (see color code on the right) showing the Z-scored gamma power signals recorded from the PrL during all 12 ESPi sessions, during five seconds before and after the beginning of long (>6 s) investigation bouts towards the isolated stimulus animal, using 0.5 s bins. Time ‘0’ mark the beginning of bout. Note the apparent reduction at the beginning of bout. **H.** As in **G**, for the group-housed stimulus. Note the apparent increase in gamma power at the beginning of bout. **I.** Super-imposed traces representing the mean (±SEM) of the signals shown in **G-H**, for both stimuli. The yellow bar marks the five minutes after the beginning of bout, across which a statistical difference was observed between the two stimuli. Unpaired t-test, n = 12 sessions, t (22) = 3.095, \*\**p* = 0.0053.

### CamK2-positive PrL neurons exhibit different stimulus-specific responses in the SP and ESPs tasks

As our results regarding PrL involvement in emotion recognition behavior are in line with previous work showing that such behavior depends on mPFC neuronal activity ^20^, we next examined the activity of the mPFC projection neurons (*CamK2a*-expressing pyramidal cells) during ESP behavior. To that end, we injected the mPFC with AAV viral vectors used to express the genetically-encoded calcium indicator GCaMP6f under the control of the *CamK2a* promoter (**Fig. 4A**) and employed fiber photometry (**Fig. 4B**) to record calcium signals (**Fig. S2A-C)** during either SP or ESPs tasks performed by male C57BL/6J mice (**Fig. 4C-D**). We used Z-score analysis to normalize and average all signals recorded during similar events (namely, at the start and end of investigation bouts towards a certain stimulus) conducted during a given session (**Fig. S2D-I**), and subsequently analyzed the normalized calcium response across all sessions of the same task (**Fig. S2J-M**).

We first analyzed calcium signals during the SP task, during which the recorded animals showed a highly significant preference for the animal stimulus, as compared to the object (**Fig. 4C**). As expected, analysis of the calcium signals recorded before the task, when the mice investigated empty chambers with no preference (**Fig. 3A-B**), revealed no difference in chamber preference (**Fig. S3C-F**). Similarly, no significant difference in calcium signals generated in response to the two stimuli was observed during short investigation bouts during the task (**Fig. S4A-C**). However, we did find a statistically significant difference in calcium signals generated in response to the two stimuli at the beginning of long investigation bouts. Specifically, we observed an excitatory response during the initial phase (i.e., in the first six seconds) of object investigation, whereas a contrasting inhibitory response occurred during the same period of social investigation events (**Fig. 4E-G**). Since we only considered investigation bouts longer than six seconds for this analysis, this difference in calcium signals between the social and object stimuli cannot be attributed to variations in bout duration between the two. Moreover, upon analyzing the signals at the end of the long investigation bouts, we noted an inverse relationship between the responses to the two stimuli (**Fig. 4H-J**), with object investigation being characterized by inhibition and social investigation being characterized by excitation (**Fig. 4J-K**). Since the change at the end of an investigation reflects the return of neural activity to baseline, these results further suggest that during the SP task, PrL pyramidal neurons are excited throughout long object investigation bouts and inhibited throughout long social investigation bouts.

**Figure 4.**
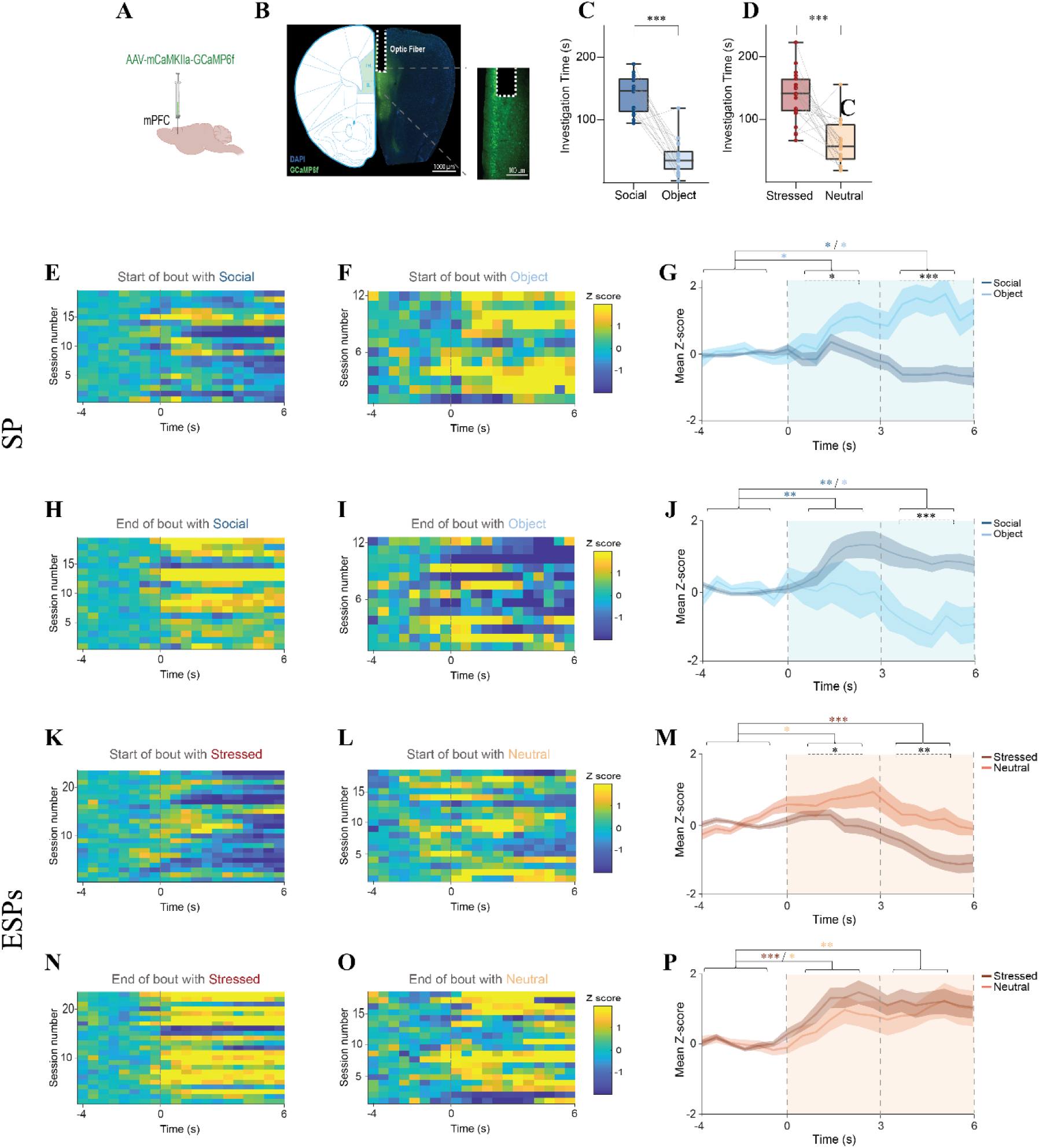
*CamK2a*-positive mPFC neurons exhibit differential responses to the stimuli in the SP task. **A.** Schematic representation of the GCaMP6-expressing AAV vector injected in the mPFC. **B.** Fluorescent microscgraph of an AAV vector-infected mPFC slice, showing GCaMP6-expression (green) and optic fiber location (dashed line). The inset shows the fiber tip at higher resolution. **C.** Mean time for investigation of the animal (blue) or object (light blue) stimulus during the SP task (n=18 sessions from 9 subjects), as measured using fiber photometry. \*\*\**p*<0.001, paired samples t-test. **D.** Mean time for investigation of stressed (brown) or neutral (light brown) stimulus animals during ESP sessions (n=23 sessions from 13 subjects), as measured using fiber photometry. \*\*\**p*<0.001, paired samples t-test. **E.** Heat-map (see color code to the right of **E**) of the Z-scored calcium signals at the beginning (first six seconds) of bout, averaged across all long investigation bouts towards the stimulus animals, shown for all sessions of the SP task (n=18 sessions, with each line corresponding to a single session), using 0.5 sec bins. Time ‘0’ represents the beginning of the bout. **F.** As in **D**, for the object stimulus in the same experiment (n=11 sessions). Note that the number of sessions was lower than in **D**, due to the lack of long bouts towards the object in some SP sessions. **G.** Super-imposed traces of the mean (±SEM) Z-scored calcium signals shown in **E-F**, averaged across all SP sessions. Dashed lines represent three distinct time windows of the event, which are used for the statistical analysis. Asterisks, representing a statistical comparison between the stimuli and across time windows, are color-coded according to the type of comparison. Blue and light blue asterisks represent a significant difference for social and object investigation, respectively, during the specific time window, relative to baseline, while black asterisks represent a difference in responses to the two stimuli during the same time window. A significant effect was found for the stimulus X time interaction (*p*<0.001) by two-way MM ANOVA. **H-J.** As in **E-G**, for the end of long bouts of investigation. A significant effect was found for the stimulus X time interaction (*p*<0.001) by two-way MM ANOVA. **K-M.** As in **E-G**, for the beginning of long investigation bouts towards the stressed (brown) or neutral (light brown) stimulus animals during the ESP task (n_stressed_=23 sessions, n_neutral_=16 sessions). A significant effect was found for the stimulus X time interaction (*p*<0.01) by two-way MM ANOVA. **N-P.** As in **K-M**, for the end of long bouts of investigation. A significant effect was only found for time (*p*<0.001) by two-way MM ANOVA. \**p*<0.05, \*\**p*<0.01, \*\*\**p*<0.001, *post-hoc* paired (within stimulus differences in time) or independent (differences between stimuli within time windows) t-test with Holm-Šídák correction for multiple comparisons following detection of main effects by ANOVA

We next analyzed calcium responses recorded during the ESPs task. As with the SP task, we found no significant difference in the responses to the two stimuli at the beginning of short investigation bouts (**Fig. S4D-F**), while a robust difference was observed when long investigation bouts were analyzed. Specifically, significant excitation was found at the beginning of investigation bouts towards neutral stimulus animals, whereas inhibition was observed at the beginning of investigations of stressed stimulus animals (**Fig. 4K-M**). Similar results were obtained with the ESPi task (**Fig S5A-E**). These results, along with those from the SP task, suggest that investigation of non-preferred stimuli elicits an excitatory response in mPFC pyramidal neurons, while investigation of preferred stimuli elicits an inhibitory response. However, when analyzing responses recorded at the end of long investigation bouts during the ESPs task, we found an increased signal at the end of investigations of both stimuli, suggesting that the cells had been previously inhibited (**Fig. 4N-P**). Similar results were found with the ESPi task (**Fig S5F-H**). We then noticed that the excitation response at the beginning of investigation events towards the neutral animal was only transient and was abolished after several seconds (**Fig. 4M**). Accordingly, when analyzing the calcium signals at the beginning of investigation bouts with a longer time window, we found that, unlike the signals recorded in the SP task, which showed persistent differences between the stimuli (Fig. **5A-D**), the responses towards both stimuli in the ESPs task reached a similar level of inhibition after about 10 s (Fig. **5E-H**). This inhibition seems to account for the similarity observed in excitation at the end of investigations of both stimuli (Fig. 5I-J). Thus, in contrast to the sustained excitation seen during investigation of the non-preferred (object) stimulus in the SP task (Fig. 6C), the excitation noted during investigation of the non-preferred (neutral animal) stimulus was transient and lasted for only a few seconds at the onset of each investigation bout in the ESPs task (Fig. 6G).

Accordingly, when directly comparing the calcium responses of mPFC pyramidal neurons at the beginning of investigation bouts in the SP and ESPs tasks, we found similar inhibitory responses towards the preferred stimuli (Fig. 6I-J). At the same time, responses towards the less preferred stimuli started to show a statistically significant difference about three seconds after the beginning of the bout (Fig. 6K-L). Overall, these results suggest that although the differential response of mPFC pyramidal neurons towards the different stimuli persisted throughout the investigation event in the SP task, the response was only differential at the beginning of investigation in the ESPs task.

**Figure 5.**
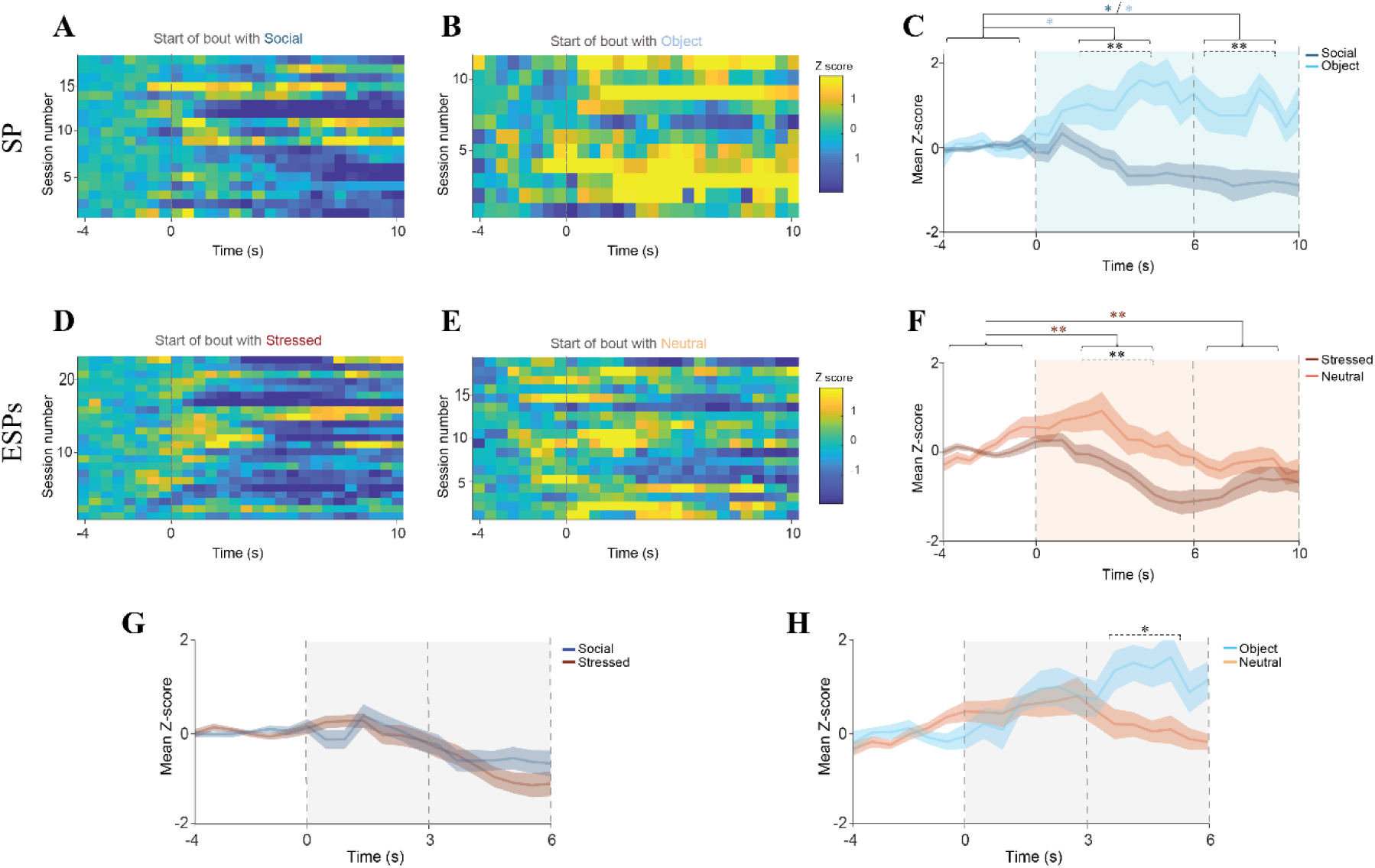
Excitatory responses of CamK2-positive mPFC neurons are persistent in the SP task yet transient in the ESPs task. **A.** Heat-map (see color code to the right of **B**) of the Z-scored calcium signals in the first 10 seconds of investigation bouts, averaged across all long bouts towards the stimulus animals, shown for all sessions of the SP task (n=18 sessions, with each line corresponding to a single session), using 0.5 s bins. Time ‘0’ represent the beginning of the bout. **B.** As in **A**, for the object stimulus in the same experiments. Note that the number of sessions is lower than in **A**, due to the lack of long bouts of investigation of the object in some sessions (n=11 sessions). **C.** Super-imposed traces of the mean (±SEM) Z-scored calcium signals shown in **A-B**, averaged across all sessions. Dashed lines represent three distinct time windows of the event, which were used for the statistical analysis. Asterisks, representing a statistical comparison between responses to stimuli and across time windows, are color-coded according to the type of comparison. Blue and light blue asterisks represent a significant difference between social and object investigations, respectively, during the specific time window, as compared to baseline, while black asterisks represent a difference between responses to the two stimuli during the same time window. A significant effect was found for the stimulus X time interaction (*p*<0.001) by two-way MM ANOVA. **D-F.** As in **A-C**, for the ESPs task (n_stressed_=23 sessions, n_neutral_=16 sessions). A significant effect was found for the stimulus X time interaction (*p*<0.001) by two-way MM ANOVA. **G.** As in **C**, calcium signals at the beginning of long investigation bouts towards the social stimulus in the SP task (blue) and the stressed stimulus in the ESPs task (Brown), shown in **C, F**. A significant effect was found for time (*p*<0.001) by two-way MM ANOVA. **H.** As in **I**, for the object in the SP task (light blue) and the stressed stimulus animal in the ESPs task (light brown). A significant effect was found for the time X stimulus interaction (*p*<0.01) by two-way MM ANOVA. \**p*<0.05, \*\**p*<0.01, \*\*\**p*<0.001, *post-hoc* paired (within stimulus differences in time) or independent (differences between stimuli within time windows) t-test with Holm-Šídák correction for multiple comparisons following determination of main effects by ANOVA.

### Optogenetic stimulation of mPFC *CamK2a*-expressing neurons specifically affects ESPs behavior

To examine if the differential responses of mPFC pyramidal neurons to the various stimuli are essential for the behavioral preference exhibited by the animals, we performed optogenetic stimulation of mPFC *CamK2a*-expressing neurons so as to interfere with their activity (Fig. 7A). To that end, we used AAV viral vectors to express the excitatory opsin channelrhodopsin 2.0 (ChR2) under control of the *CamK2a* promoter in the mPFC (**Fig. 6A**), followed by optic fiber implantation for blue light delivery to the mPFC (**Fig. 6B**). Optogenetic stimulation during investigation of one of the empty chambers before the task did not create any behavioral preference for a given chamber (**Fig. 6C**). When we excited mPFC pyramidal neurons during the SP task by applying photo-stimulation (15 ms pulses at 10 Hz) during the first second of investigation bouts towards either the animal (social) or object stimuli (**Fig. 6D-E**), no change in the social preference of the subjects for either stimulus was seen, as compared to no stimulation controls (**Fig. 6F**). Moreover, no changes in motor activity variables, such as number of transitions between stimuli and traveled distance, were seen in either protocol (**Fig. S6A-C**). As a positive control, we applied free stimulations of 10 Hz across the entire encounter period of the SP task and found, in agreement with previous studies ^30,31^, that such stimulation abolished social preference of the subjects (**Fig. 6D**). However, when we applied the same set of stimulation protocols during the ESPs task (**Fig. 6H-I**), we observed very different results. Firstly, stimulating during the first second of investigation bouts towards the stressed animals (i.e, the preferred stimulus) abolished behavioral preference. Secondly, when we applied the stimulation during the first second of investigation bouts towards the neutral stimulus animals (i.e., the non-preferred stimulus), subject preference to investigate the stressed animal was augmented and grew even stronger than did their preference without any stimulation (**Fig. 6J**). Notably, these stimulation protocols did not cause any change in motor activity variables (**Fig. S6D-E**). Furthermore, there was no difference in the mean number of stimuli applied during both stimulation protocols, thus excluding the possibility that such a difference was the source of the differential response seen (**Fig. S6F**). Similar to the SP task, free stimulation applied during the ESPs task abolished the preference of the mice to investigate one stimulus more than the other (**Fig. 6K**). Altogether, these results suggest that the subject’s preference to investigate a stressed conspecific more than a neutral one in the ESPs task depends on the transiently differential response of mPFC pyramidal neurons to distinct stimulus animals. In contrast, the SP task seems to be resilient to a brief stimulation applied at the beginning of investigation bouts, most probably because the differential response seen during the SP task is more persistent than is that seen in the ESPs task.

**Figure 6.**
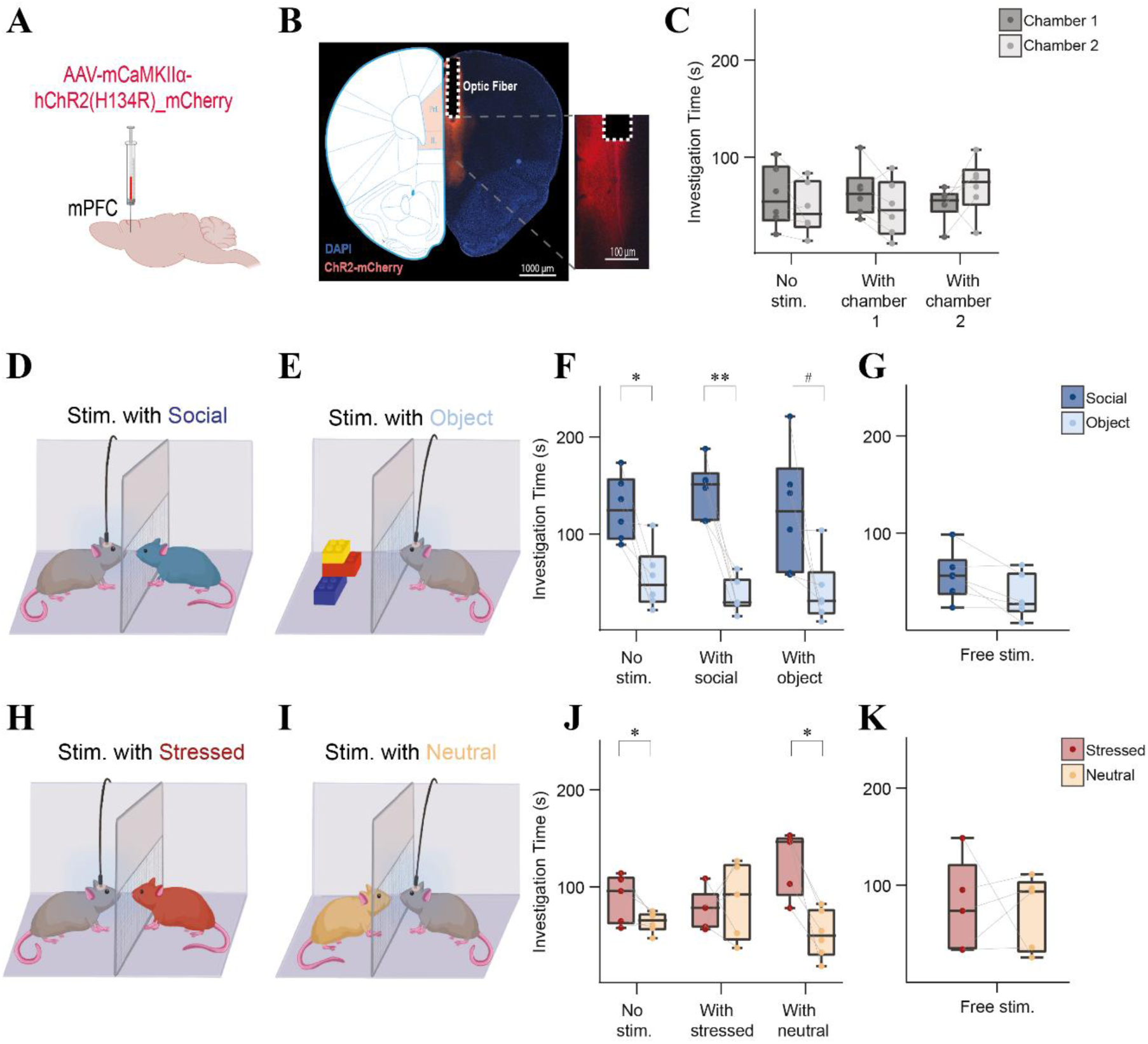
Optogenetic stimulation of CamK2a-positive mPFC neurons has a differential effect in the SP and ESPs tasks. **A.** Schematic representation of ChR2-expressing AAV vector injection in the mPFC. **B.** Fluorescent micrograph of an infected mPFC slice, showing ChR2-mCherry expression (red) and the position of the optic fiber (dashed line). The inset shows the location of the fiber tip at higher resolution. **C.** Mean investigation time of each chamber during a five min exposure to two empty chambers (n=6 sessions), with no stimulation or when optogenetic stimulation was consistently applied for one second at the beginning of events of investigation of chamber 1 (grey) or chamber 2 (light grey). **D.** Schematic representation of the ‘with social’ stimulation protocol, where optogenetic stimulation was applied in the first second of social investigation bouts during the SP task. **E.** As in **D**, for object investigation bouts (‘with object’). **F.** Mean time of investigation of the social (blue) and object (light blue) stimulus during the SP task (n=6 sessions), while applying each of the three stimulation protocols denoted below the bars. A significant effect was found for stimulus (*p*<0.001) by two-way MM ANOVA. **G.** As in **F**, when a free optogenetic stimulation of 10 Hz was applied across the five min SP task. Note the reduction in social investigation to the same level as seen with object investigation in this case. **H.** Schematic representation of the ‘with stressed stimulation protocol, where optogenetic stimulation was applied at the first second of investigation bouts towards the stressed stimulus animal during the ESPs task. **I.** As in **H**, for investigation bouts towards the neutral stimulus animal (‘with neutral). **J.** Mean time of investigation of the stressed (red) and neutral (cream) stimulus animals during the ESPs task (n=6 sessions), using the three stimulation protocols denoted below the bars. A significant effect was found for the stimulus X condition interaction (p<0.05) by two-way MM ANOVA. **K.** As in **J**, when a free optogenetic stimulation of 10 Hz was applied across the five min session of the ESPs task. #*p*<0.07, \**p*<0.05, \*\**p*<0.01, \*\*\**p*<0.001, *post-hoc* paired t-test with Holm-Šídák correction for multiple comparisons following detection of a main effect by ANOVA.

### Prefrontal cortex neuronal activity precedes investigation behavior and predicts the type of investigated stimulus

Thus far, we demonstrated that the differential neural activity of mPFC pyramidal neurons which takes place upon investigation of stimulus animals is required for emotion recognition by male mice. Yet, this activity may participate in maintaining investigative behavior once initiated, or be involved in the decision made by a subject to start or to stop stimulus investigation. The latter possibility requires mPFC neurons to change their firing activity before the beginning or end of the investigative behavior. To differentiate between these two possibilities, we used chronically-implanted Neuropixels 1.0 probes (**Fig. 7A-B**) to record neuronal activity at the single-cell level in the prefrontal cortex of subject mice conducting the SP (**Fig. S7A**), ESPs (**Fig. 7C**) and ESPi (**Fig. S7C**) tasks, at the high temporal resolution scale of milliseconds. We analyzed the activity of >1000 units per task (1009 for ESPs, 1342 for ESPi and 1044 for SP), recorded from the mPFC (PrL and IL) and anterior cingulate cortex (ACC) of four subject mice performing each task 2-4 times (sessions), with each session being conducted on different days and considered separately. When analyzing the firing frequency of each unit during the five min pre-encounter period, we found that in all tasks, more than 80% of the units fired below 10 Hz (**Fig. 7D** and **Fig. S7B, D**), while more than 50% of them fired below 5 Hz. These results are in accordance with a previous study that used a similar recording technique and reported about 90% putative excitatory neurons among the units recorded ^32^.

We used Z-score analysis to assess the activity of each unit in synchrony with either the beginning or end of long (>6 s) investigation bouts, separately for each stimulus in each task (see examples in **Fig. 7E-J**). As is apparent from **Fig. 7K-L**, showing results from mPFC units recorded during ESPs sessions, with the activity of each neuron being averaged across all investigation bouts towards a given stimulus in each session, we found a spectrum of responses ranging from strong excitation to strong inhibition at both the beginning and end of such investigations of each stimulus. When averaging the Z-score traces of all units, we observed a mean excitatory response at both the beginning and end of investigation bouts towards both stimulus animals, which seemed to begin about one second before the behavioral event itself. We also observed significantly higher excitation towards the neutral stimulus animal, as opposed to the stressed animal, specifically 0.5 s before the beginning and 1.5 s after the end of the investigation bouts (**Fig. 7M-N**). Similar results were found for ESPi and SP sessions, albeit with no significant difference between the responses to stimuli (**Fig. S7E-L**).

**Figure 7.**
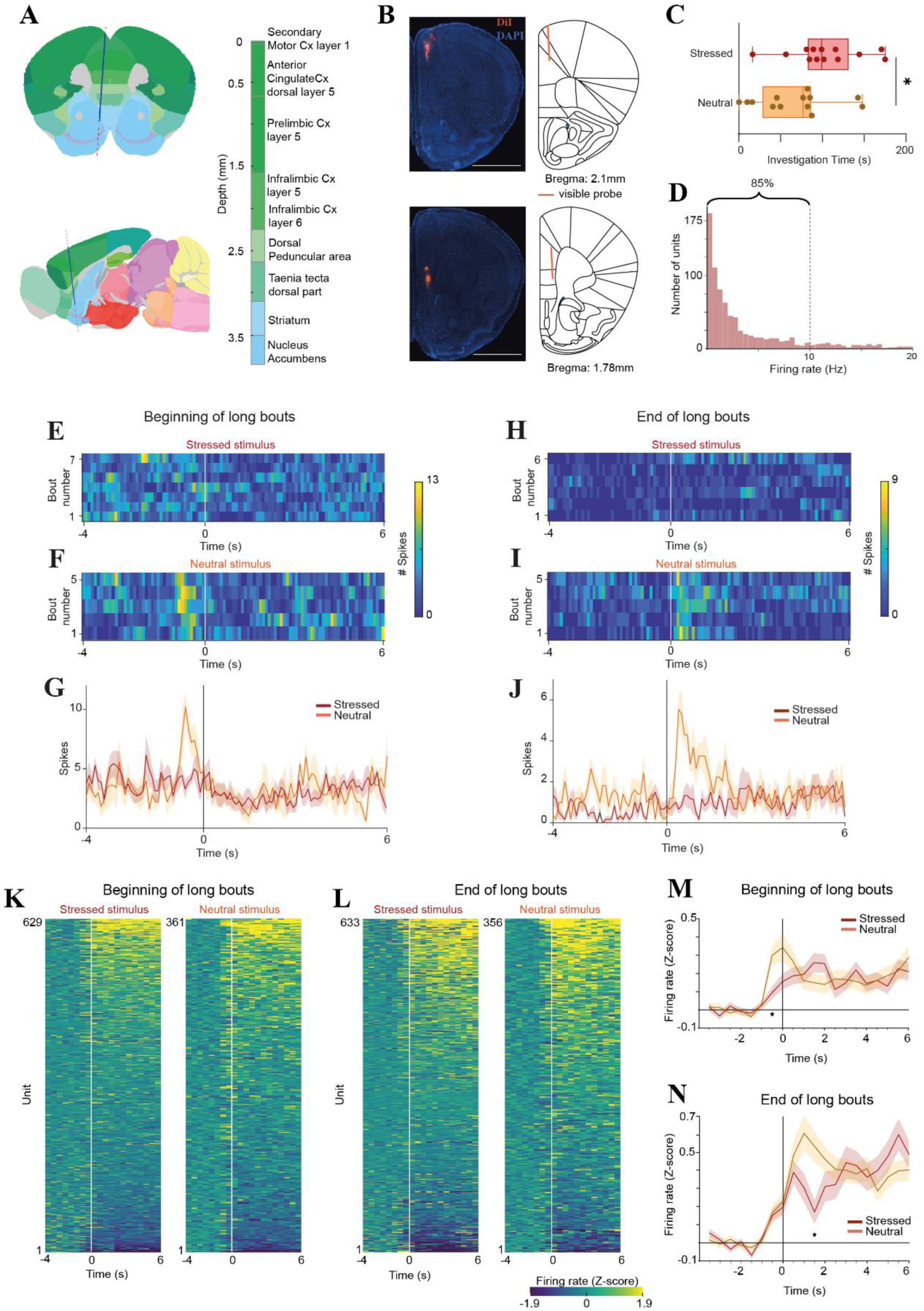
Differential response of prefrontal cortex single units to the distinct stimuli during the ESPs task. **A.** Top panel - schematic representation of a coronal forebrain slice showing the trajectory of the implanted Neuropixels 1.0 probe, with the various brain regions color-coded according to their depth. See the color code on the right. Bottom panel – the same, for a sagittal slice. **B.** Left panel - two fluorescent micrographs of consecutive DAPI-stained slices, with the trajectory of the DiI-labeled probe marked in red in the diagram on the left. Right panel - the corresponding schematic brain atlas representations. **C.** Median time dedicated by recorded male subjects for investigation of stressed (red) and neutral (orange) stimulus animals during the ESPs task (n=13 sessions, performed by 4 subjects). **D.** Distribution of the of the number of recorded units according to r firing frequency during the baseline (pre-encounter) period (0.5 Hz bins). Note that 85% of the units fired below 10 Hz. **E.** A heat-map of neural activity (using 100 ms bins) recorded from a single PrL unit four seconds before and six seconds after the start (time ‘0’) of each long bout of investigation of a stressed stimulus animal during a single ESPs session. Each line represents a single bout. **F.** As in **E**, for the neutral stimulus animal in the same session. **G.** Super-imposed traces of the mean neural activity shown in E and F. Note the strong excitation just before the beginning of investigation bouts towards the neutral stimulus. **H-J.** As in **E-G**, for the end of bouts in the same session. Note the strong excitation just after the end of bouts towards the neutral stimulus animals. **K.** Heat-maps of the mean Z-scored firing rate recorded at the beginning of long investigation bouts towards the stressed (left) and neutral (right) stimulus animals, for all units that fired at least one spike during this period (hence, the distinct number of units per stimulus animal). The various units are arranged according to their responses from strongest excitation (top) to strongest inhibition (bottom), separately for each stimulus animal. **L.** As in **K**, for the end of bout. **M.** Super-imposed traces of the mean Z-scored firing rate of all neurons shown in **K**. Note the significantly stronger excitation during investigation bouts towards the neutral stimulus, specifically just before the beginning of the bout. Multiple unpaired t-test (each 0.5-s bin), time bin (-0.5 s), BHFDR corrected, \**p* = 0.0102. **N.** Super-imposed traces of the mean Z-scored firing rate of all neurons shown in **L**. Note the significantly stronger excitation during investigation bouts towards the neutral stimulus, specifically just after the beginning of the bout. Multiple unpaired t-test (each 0.5 s bin), time bin (1.5 s), BHFDR corrected, \**p* = 0.0131.

We next analyzed the types of responses at the beginning and end of all investigation bouts towards each stimulus, across all units. To that end, we used a criterion of mean Z-score (across the bout) of ±1.9 as defining a responsive unit. In all three tasks, we found that <20% of the cells responded with excitation to at least one of the four behavioral events. Most of these neurons (14-16%) responded in one event only, while <4% responded in more than one event. Similarly, we found that <10% of the units responded with inhibition, with most of them (6-9%) responding in one event alone and <2% responding in more than one event (**Figure S8**). Together, these observations suggest that most responding units are event-specific.

To analyze the data at the population level, we used hierarchical clustering to categorize the various units into functional groups, based on their responses both at the beginning and end of all investigation bouts towards each stimulus animal during each task. For the ESPs task, we counted seven distinct groups, six of which comprised more than 30 neurons and hence, were further considered (**Fig. 8A**). Most of these neurons were recorded from layer 5 of the PrL and ACC (**Fig. 8B**). Group 5 comprised about half of the neurons (55.9%; **Fig. 8H**) and, as compared to the baseline, did not show any significant response in any of the events (**Fig. 8H**). The other five groups (**Fig. 8D-G, I**) responded significantly to at least one of the events, displaying either excitation or inhibition. In all cases, significant excitatory responses were specific to a single stimulus. For example, Group 1 responded with strong excitation at the beginning of investigation bouts towards the neutral stimulus animal, while group 2 responded with strong excitation at the beginning of investigation bouts towards the stressed stimulus. Similarly, Group 4 responded with excitation at the end of investigation bouts towards the neutral stimulus animals, while Group 7 responded with excitation at the end of investigation bouts towards the stressed stimulus. Finally, Group 2 and Group 4 responded in opposite direction at the beginning and end of investigation bouts towards the same stimulus. It should be noted that in most cases, responses had already begun one second before the beginning of the behavioral event (**Fig. 8D-F, I**). Qualitatively similar results were found for the SP (**Fig. S9**) and ESpi (**Fig. S10**) tasks.

**Figure 8.**
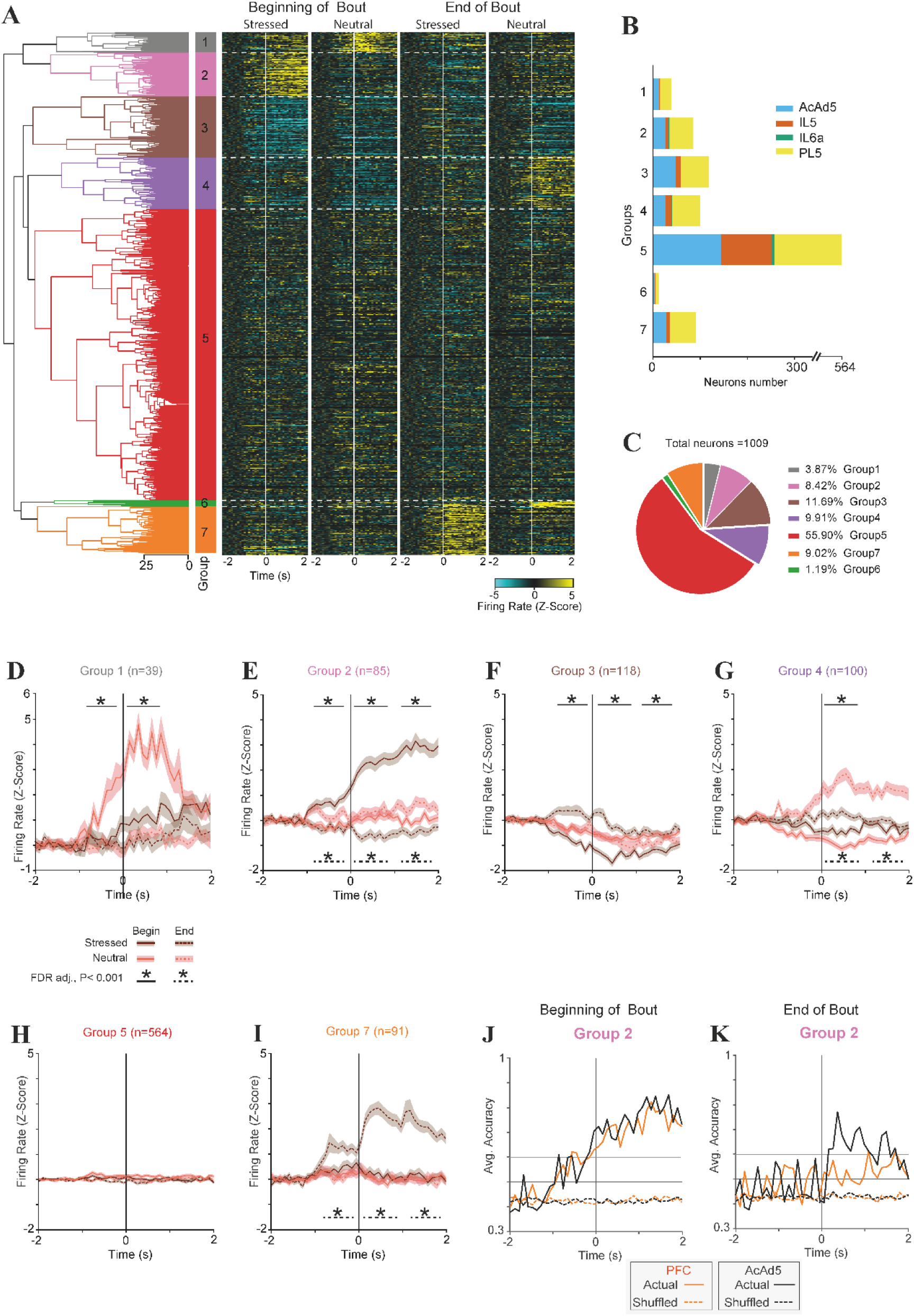
Distinct groups of prefrontal neurons respond differentially to distinct stimuli during the ESPs task. **A.** A dendrogram of a hierarchical clustering of all cells (1009) that fired during either the beginning or end of investigation bouts towards any of the stimuli during the ESPs task, according to their Z-scored response. A heat-map showing the mean firing response of each cell during all four event types (denoted above) is displayed to the right of the dendrogram. **B.** Color-coded distribution of the units comprising each group to four recorded prefrontal regions, as detailed in the legend on the right: AcdAd5 – layer 5 of the dorsal anterior cingulate cortex, IL5 – layer 5 of the infralimbic cortex, IL6a - layer 6a of the infralimbic cortex, and PL5 – layer 5 of the prelimbic cortex. Note that most of the responding units are from the PrL area, while the IL and ACC are over-represented in the non-responding units (group 5). **C.** A color-coded pie chart showing the fraction (in %) of each group among all recorded units. **D.** Super-imposed traces of the mean Z-scored firing rate of the units included in Group 1 at the beginning (solid lines) and end (dashes lines) of investigation bouts towards the neutral (red) or stressed (brown) stimuli. \**p*<0.001 in FDR-adjusted *post-hoc* t-test between the responses (averaged at 1-s bins) at the beginning or end of investigation bouts for the two stimuli, following two-way RM ANOVA analysis of time bins and stimulus. Note that the significant difference starts as early as one second before the investigation bout. **E.** As in **D**, For Group 2. Here, we found a significant difference between stimuli both at the beginning (solid lines, above) and end (dashed lines, below) of investigation bouts. Again, the significant difference started before the behavioral event. **F-I.** As in **D**, for Groups 3, 4, 6 and 7, respectively. **J.** Traces representing the accuracy of a logistic regression model predicting the identity of the investigated stimulus according to the firing rate of Group 2 neurons at the beginning of investigation bouts, as shown in **E**, analyzed separately for mPFC (orange trace) and ACC (black trace) neurons. The results of the same model using shuffled data are shown as dashed lines, using the same colors. Note that the predictive accuracy started to rise about 1 s before the beginning of the bout, for both PFC and ACC neurons. **K.** As in **J**, for the same neurons during the end of bout. Note that in this case, ACC neurons reached an accuracy >70%, while mPFC neuronal accuracy was <60%.

Finally, we examined if the differential responses of each neuronal group at either the beginning or end of investigation bouts could be used to distinguish between the two stimuli. To that end, we applied a linear logistic regression classifier to the firing rate of each group and examined, at each time bin, classifier accuracy in predicting the investigated stimulus, separately for mPFC (PrL and IL) or ACC neurons. As exemplified in **Fig. 8J**, for group 2 at the beginning of bout, (see all responding groups of the SP and ESPs tasks in **Fig. S11**), in many cases, the model achieved an accuracy of >70% in predicting the investigated stimulus, both for the SP and ESPs tasks. As a control, we used the model on shuffled data from each group, which always yielded a low accuracy of 40-45%. Moreover, in most of these cases, both the mPFC and ACC performed similarly to each other, with accuracy reaching >60% even before the beginning of behavioral events (For examples, **Fig. S11B, J, R, N, T**). However, in some cases, one of the brain regions showed higher accuracy than did the other (see for example **Fig. 8K** and Fig. **S11H**), suggesting that there might be a difference between the two brain regions (i.e, the ACC and mPFC).

Overall, these results suggest that the differential responses of mPFC neurons are not only essential for murine emotion recognition but are also involved in decisions made by the subject as to whether to start or stop social investigation behavior. Thus, these neurons can be assigned a role in social decision making.

## Discussion

Emotion recognition, i.e., the ability to assess the emotional state of others, is crucial for prosocial behavior and empathy ^21^. Here, we demonstrated that male and female mice discriminate between same-sex conspecifics on the basis of their emotional state. Moreover, while both male and female subjects exhibited relief- and stress-state preference, only males preferred isolated over group-housed conspecifics. In contrast, female did not show such preference, even when males served as stimuli, thus excluding the possibility that the difference between males and females is caused by the sex of the stimulus animals. The sex-dependent nature of ESPi may be due to an inability of female subjects to recognize a state of isolation in the stimulus animals or due to their indifference towards group-housed and isolated conspecifics, an issue requiring future study. Finally, we showed that at least male mice can distinguish between various emotional states (namely, relief, stress and social isolation). This discrimination does not necessarily mean that males qualitatively differentiate between the distinct states. Instead, subjects may be able to distinguish distinct states by the level of arousal they elicit. Future studies using varying levels of each state may be required to address this issue.

Since emotion recognition is known to be impaired in human individuals diagnosed with ASD ^9–11^, we reasoned that ASD mouse models may be prone to impairments in this behavior. Using a battery of social discrimination tasks, we found that *Shank3*-KO male mice exhibit specific deficits in the SNP and ESPs tasks, while performing normally in the SP and SxP tasks. It should be noted that due to large differences in the experimental arenas, our SP and SNP tasks are not identical to the sociability and SNP tasks used in the three-chamber test. Therefore, it is not surprising that our results from the SP task differ from sociability results reported by a previous study, which used the three-chamber test with the same mouse line ^27^. Nevertheless, both studies found similar deficits in the SNP task. Importantly, several established mouse models of ASD were shown to perform normally on our version of the SP task, while all models showed deficits in various versions of the ESP task ^19,28^. These results demonstrate the robustness of social preference behavior of mice and suggest that it is much more resilient to disturbances than is emotion recognition. This difference in resilience between the SP and ESP tasks, further confirmed by our optogenetic stimulation experiments, is likely due to the various level of challenge they impose on a subject. While social preference behavior relies on the extensive differences between object and social stimuli, i.e., animals, distinguishing between animals with distinct emotional states most probably reflects subtle differences in their odor, vocalization or motor behavior. Therefore, emotion recognition is expected to be more challenging that social preference and hence, more sensitive to external or internal disturbances. This may explain the common impairment in emotion recognition exhibited by ASD individuals ^9,11^. Altogether, our results support the notion that the ESP paradigm is a valid behavioral model for human emotion recognition, as previously suggested by others ^21^. Thus, this paradigm can be employed to unravel brain mechanisms and circuitries underlying emotion recognition in rodents, leading to new insight into similar mechanisms in the human brain that subserve human emotion recognition and prosocial behavior.

To locate brain regions involved in murine ESP behavior, we used chronically-implanted microelectrode arrays in behaving mice to record LFP signals from multiple brain regions involved in social behavior ^33^. We reasoned that brain regions involved in discrimination between aroused and non-aroused conspecifics will show differences in their neural activity during interactions with the two types of stimulus animals. Of the eight brain regions we have analyzed in this study, only the PrL showed such a difference in rhythmic LFP signals. Since the mPFC was previously reported to be involved in ESP ^20^ and given our recent findings demonstrating that the level of synchronization in theta and gamma rhythmicity between the PrL and the hypothalamic paraventricular nucleus (PVN) is significantly modified during social behavior in another murine model for ASD (*Cntnap2*-KO mice) ^28^, we further explore the ESP-related neuronal activity in this region at both the population (using fiber photometry recordings) and single-cell (using Neuropixels recordings) levels. Using AAV vector-mediated gene delivery for fiber photometry allowed us to specifically target the PrL pyramidal neurons, using the *CamK2a* promoter to induce expression of GCaMP6f. These excitatory neurons are known to project to many brain areas ^33^, where they presumably regulate distinct aspects of behavior. However, a limitation of our study is that we did not distinguish between the various sub-populations of PrL pyramidal neurons. Still, it should be noted that a recent study which used fiber photometry to explore the responses of PrL neurons projecting to either the contralateral PrL, nucleus accumbens, or ventral tegmental area using various paradigms of avoidance behavior observed qualitatively similar responses in all sub-populations ^34^. Our fiber photometry recordings clearly demonstrated a significant difference in calcium signals recorded at the beginning of long investigation bouts towards preferred or non-preferred stimuli in the SP, ESPi and ESPs tasks. Interestingly, in all tasks, we observed excitation at the beginning of investigation bouts towards the non-preferred stimulus, while inhibition was recorded during investigation of the preferred stimulus. Our finding that these differences could be observed during long investigation bouts, where clear behavioral preference is observed, but not during short bouts where we found no preference, further supports association between differential neuronal responses and behavioral preference.

The differential responses of PrL pyramidal neurons at the beginning of long investigation bouts towards the preferred or non-preferred stimuli is in accordance with the well-known role of the mPFC in controlling impulses ^35^ and of the PrL area, in particular, in regulating avoidance behavior ^36,37^. Moreover, the differential responses towards social and object stimuli that we observed during the SP task resemble the responses reported in IL pyramidal neurons to social and object stimuli in a previous study using the same recording technique ^38^. Yet, we observed a clear distinction between the SP and ESPs tasks in the form of differential PrL responses. While in the SP task the differential response persisted throughout the investigation bout, in the ESPs task, the differential response was more transient in nature. Accordingly, the excitatory response during investigation bouts towards the neutral stimulus animal turned inhibitory after about three seconds, thus becoming similar to the response towards the stressed stimulus animal. Accordingly, we could not observe any difference in the responses to the two stimuli at the end of investigation bouts. Such a dynamic change in the differential response across the duration of investigation may be dictated by various inputs from the ventral hippocampus to the PrL, which were recently shown to be modulated in a similar manner and to control approach/avoidance behavior ^39^. Alternatively, our observations may reflect dynamic change in the behavior of the neutral stimulus animal during interaction with the subject. This possibility, however, seems unlikely, as the neutral stimulus animal used in the ESPs task is the same type of animal used as a social stimulus in the SP task. Regardless of its cause, we suggest that the transient nature of the differential response towards the stimuli in the ESPs task, as compared to the SP task, makes this task more sensitive to internal or external disturbances, as demonstrated by the higher sensitivity of the ESPs task to optogenetic stimulation.

Our finding that free stimulation at 10 Hz throughout the SP task abolished subject preference for the social over the object stimulus by reducing subject motivation for social investigation is not surprising, given that similar results were previous reported by multiple studies ^30,31^. This result was interpreted as reflecting a social avoidance state induced by intensive PrL stimulation. However, in accordance with a previous study ^30^, we found that stimulating PrL pyramidal neurons at the beginning of investigation bouts in the SP task did not cause any noticeable behavioral change. Such an investigation bout-specific stimulation was much weaker (<10% of stimuli) than what was seen with the free stimulation protocol, and hence may not induce a social avoidance state, as does free stimulation. In contrast, in the ESPs task, the same investigation bout-specific stimulation protocols yielded a significant change in subject behavior. Our interpretation of these results is that mouse behavior in the ESPs task depends on the transiently differential response of PrL pyramidal neurons at the beginning of long investigation bouts towards the distinct stimulus animals, such that stimulation that facilitates this differential activity causes enhancement of the behavioral preference, while stimulation that abolished it eliminates the behavioral preference. The reason we did not observe similar results in the SP task is that the differential activity in this task persisted throughout investigation bouts, rather than being transient, and was thus not affected by the short (1 sec-long) stimuli we applied at the beginning of each investigation bout. It should be noted that using a closed-loop optogenetic stimulation to apply excitatory stimulation to PrL pyramidal neurons throughout bouts of social investigation, a previous study did manage to abolish the social preference of mice, in accordance with our suggestion ^30^.

Since multiple studies have implicated mPFC neuronal activity in decision-making processes in both social ^40^ and non-social ^41–43^ contexts, we explored the dynamics of mPFC neuronal activity during the SP and ESPs tasks at the single-cell level. Notably, these recordings could not distinguish between the activity of pyramidal neurons from that of interneurons. Therefore, the results of such recordings are expected to differ from those collected by fiber photometry, which were specifically obtained from *CamK2a*-expressing neurons. Nevertheless, we found that during the ESPs task, but not the SP task, the mean response to the two stimuli across all neurons differed significantly. This difference was observed in specific time windows, with higher firing being seen immediately before the beginning and after the end of long investigation bouts towards the non-preferred neutral stimulus, as compared to the preferred stressed stimulus. The fact that we did not observe such a difference in the SP and ESPi tasks suggests that it may be context-specific.

We used two distinct methods to categorize the recorded units according to their Z-scored responses. First, we used a cutoff of ±1.9 to identify responsive units in four time windows, namely, at the beginning and end of investigation bouts towards each of the two stimuli. Second, we used unsupervised hierarchical clustering to categorize the units into groups according to their response profiles in the same four time windows. Both methods revealed that in all three tasks (i.e., SP, ESPi and ESPs), excitatory responses of the recorded units were mainly specific to a single time window. Thus, it seems as if each neuronal group is involved in encoding either the beginning or end of investigation of either the preferred or non-preferred stimulus during a social discrimination task. These results are in accordance with previous studies demonstrating the involvement of PrL neurons in regulating goal-oriented behavior ^37^. Importantly, we found that in many of the neuronal groups, excitatory responses started even before the investigative behavior, suggesting that these neurons are involved in the process of taking the decision whether to start or stop investigating a certain stimulus. These results are in accordance with previous studies showing similar dynamics of PrL neuronal response relative to behavioral events, in either social ^44^ or non-social ^42^ contexts.

To summarize, or results demonstrate that mPFC differential neuronal response to conspecifics with distinct emotional state subserves the emotion recognition ability of mice, which is consistently impaired in multiple ASD mouse models. Therefore, atypical activity of these neurons, which seem to be involved in the decision whether to engage or avoid a certain conspecific in a social context, may be key to various deficits in social behavior, including emotion-recognition abilities, exhibited by individuals with ASD ^9–11^.

## Supporting information

All processed data used for generating the figures

All results of statistical tests

## Author contributions

A.N.M.: Formal analysis, Investigation, Methodology, Validation, Visualization, Writing - original draft, and Writing - review & editing; R.J.: Formal analysis, Investigation, Methodology, Validation, Visualization; N.R.; Investigation, Formal analysis; S.N.: Data curation, Project administration, Software, Validation, Visualization, Writing - original draft, and Writing - review & editing. S.W.: Conceptualization, Funding acquisition, Project administration, Resources, Supervision, Writing - original draft, and Writing - review & editing.

## Acknowledgments

We thank Boris Shklyar, Head of the Bio-imaging Unit and Eng. Alex Bizer, the experimental systems engineer of the Faculty of Natural Sciences of the University of Haifa, for their technical help. We also thank Sarah Sheikh for helping with the graphical aspects of the figures.

## Methods

### Resource availability

Further information and requests for resources and reagents should be directed to and will be fulfilled by Shlomo Wagner.

### Materials availability

This study did not generate new unique reagents.

### Data and code availability

All the data generated from experiments and the custom codes used to evaluate the conclusions of this paper would be uploaded onto a public repository upon acceptance of the paper.

## Experimental model and subject details

### Animals

#### C57BL/6J

C57BL/6J naïve adult (8-18 week-old) male and female mice (Envigo RMS, Israel) were used either as subjects or stimuli.

#### ICR

ICR (CD-1) subjects were also commercially obtained adult (8-18 week-old) male and female mice. Social stimuli for experiments with ICR or *Shank3* mutant mice were ICR adult male mice (8-18 week-old) in the ESP tasks, or juvenile ICR male mice (3-6 week-old) in the SP and SNP tasks. *<u>Shank3</u>*: B6.129-*Shank3^tm2Gfng^*/J mutant mice (*Shank3B*) ^45^ were crossed by us for five generations with ICR mice to create *Shank3B* mutant mice with a B6;ICR background. All *Shank3B*^−/−^ (KO) and wild-type (WT) mice used in this study were obtained by crossing male and female *Shank3B*^+/−^ mice born with the expected Mendelian frequencies. *Shank3B* subjects with KO or WT genotypes were used for behavioral testing at 8-18 weeks of age.

#### Genotyping

Ear tissue samples were collected from offspring mice at 21 days of age for genotyping by polymerase chain reaction (PCR) using the following primers:

WT forward primer: GAGCTCTACTCCCTTAGGACTT;

WT reverse primer: TCCCCCTTTCACTGGACACCC, yielding a 250 bp band;

Mutant forward primer: GAGCTCTACTCCCTTAGGACTT;

Mutant reverse primer: TCAGGGTTATTGTCTCATGAGC, yielding a 330 bp band.

#### Maintenance

Isolated stimuli for the ESPi task were individually housed for 1-2 weeks before an experiment. All other animals were kept in groups of 2-5 sex-matched mice per cage. All animals were housed at the animal facility of the University of Haifa under veterinary supervision, in a 12 h light/12 h dark cycle (lights on at 9 PM), with *ad libitum* access to food (standard chow diet, Envigo RMS, Israel) and water.

### Institutional review board

All experiments were performed according to the National Institutes of Health guide for the care and use of laboratory animals and approved by the Institutional Animal Care and Use Committee of the University of Haifa.

## Method details

### Experiments

Behavioral experiments were conducted in the dark phase of the dark/light cycle in a sound- and electromagnetic noise-attenuated chamber, under dim red light.

### Experimental setups

The experimental setup ^46^ consisted of a white Plexiglas arena (37 X 22 X 35 cm or 40.5 X 40.5 X 35 cm) for the 3-stimuli ESP experiments, placed in the middle of an acoustic cabinet (60 X 65 X 80 cm). Two (or three in the case of the three-stimuli ESP experiment) Plexiglas triangular chambers (12 cm isosceles, 35 cm height), into which an animal or object (plastic toy) stimulus could be introduced, were placed in two randomly selected opposite corners of the arena. A metal mesh (12 X 6 cm, 1 X 1 cm holes) placed at the bottom of the triangular chamber allowed direct interaction with the stimulus through the mesh. A high-quality monochromatic camera (Flea3 USB3, Flir) equipped with a wide-angle lens was placed at the top of the acoustic chamber and connected to a computer, enabling a clear view and recording (∼30 frames/s) of subject behavior using commercial software (FlyCapture2, FLIR).

### Behavioral paradigms

#### Social preference and social novelty preference

Subjects were taken from their home cage and placed in an arena with empty chambers for a 15 min habituation period. Throughout this time, social stimuli were placed in their chambers near the acoustic cabinet for acclimation. After habituation, the chamber containing the social and object stimuli were diagonally placed at opposite ends of the arena in a random fashion, and the social preference (SP) task was conducted for 5 min. Following the SP task, the chambers with the stimuli were removed from the arena, and the subject was left alone for 15 min. Then, the chambers were re-inserted, this time at the other two corners of the arena, with one containing the same social stimulus used for the SP task (familiar stimulus) and the other containing a novel stimulus. The social novelty (SNP) task lasted 5 min.

#### Sex Preference

The sex preference (SxP) task consisted of 15 min habituation to the arena with empty chambers, followed by exposing the subject to both adult male and female social stimuli located in individual chambers found at opposite corners of the arena for 5 min.

#### ESP tasks

*ESPr* - The task consists of 15 min of habituation followed by a 5 min period in which the subject mouse was introduced to a neutral stimulus animal and to another stimulus animal that had received 1 h *ad libitum* access to water after 23 h of water deprivation. Each “relieved” stimulus animal was used as a stimulus for only two consecutive sessions.

*ESPs* - The task consists of 15 min of habituation followed by a 5 min period in which the subject mouse was introduced to a neutral stimulus animal and to another stimulus animal that had been constrained for 15 min in a 50 ml tube pierced with multiple holes for ventilation. Each “stressed” stimulus animal was used as a stimulus for only two consecutive sessions.

*ESPi* - The task consists of 15 min of habituation followed by a 5 min period in which the subject mouse was introduced to a group-housed stimulus animal, and a socially isolated (1-2 weeks) stimulus animal. Each isolated stimulus was used for two non-consecutive tests.

### Stereotactic surgery for viral injection and optic fiber or electrode array implantation

#### Surgery

Mice were anesthetized and analagized with an intraperitoneal injection of a mixture of ketamine, domitor, and the painkiller norocarp (0.13 mg/g, 0.01 mg/g, and 0.005 mg/g, respectively). Anesthesia levels were monitored by testing toe pinch reflexes and maintained using isoflurane (0.2%; SomnoSuite, Kent Scientific Corporation). Body temperature was kept constant at approximately 37°C using a closed-loop custom-made temperature controller connected to a temperature probe and a heating pad placed under the animal. Ophthalmic ointment (Duratears, Alcon, Couvreur, NV) was applied to maintain eye moisture. Anesthetized animals were fixed in a stereotaxic apparatus (Kopf), with the head flat. The head was shaved, the scalp was disinfected, and the skin was removed to expose the skull. The skull was then leveled using bregma–lambda measurements.

#### Optogenetics and fiber photometry

The region of interest was marked and a hole (unilateral, right hemisphere) was drilled for viral injection and optic fiber implantation. Additional holes were drilled for supporting screws. Viral injection was performed using a glass capillary filled with the virus (ssAAV-1/2-mCaMK2a-GCaMP6f-WPRE-SV40p(A) for fiber photometry recordings or ssAAV-1/2-mCaMK2α-hChR2(H134R)_mCherry-WPRE-SV40p(A) for optogenetic stimulation; Zurich VVF, Switzerland) was slowly lowered into the mPFC (AP,+1.9 mm; ML, -0.35 mm; DV, +2.2 mm) and left in place for 5 min prior to injection and 10 min following injection to prevent retraction of the virus. A total of 300 nl of the virus was then delivered by manual application using a 50 ml syringe connected to the glass capillary (BRAND, disposable BLAUBRAND micropipettes, intraMark, 5*μl*). Following viral injection, an optic fiber (Doric lenses, 200 μm, NA 0.66, 3 mm long, zirconia ferrule, flat-fiber tip) was inserted into the mPFC with the same coordinates as the viral injection, and placed 200 μm above the injection depth. Screws and optic fiber were fixed to the skull plate using dental cement (UNIFAST, GC America).

#### Multi-electrode arrays (EAr) implantation

Following skull alignment, two burr holes were drilled to allow placement of ground and reference wires (silver wire, 127 µm, 300-500 Ω; A-M Systems, Carlsborg, WA). Two watch screws (0-80, 1/16”, M1.4) were inserted into the temporal bone to support the EAr with dental cement. Four points (at coordinates: AP = 2 mm, ML= -0.3 mm; AP = 1 mm, ML= -2.3 mm; AP = -2 mm, ML= -2.3 mm; AP = -2 mm, ML= -0.3 mm) were marked over the left hemisphere with a marker. The skull covering these marked coordinates was removed after smoothening the bone with a dental drill, and the exposed brain was kept moist with cold, sterile saline. We custom-designed the EAr ^47^ from 8 to 12 individual 50 µm formvar-insulated tungsten wires (50-150 kΩ, #CFW2032882; California Wire Company) to target the PrL, IL, AcbC, AcbSh, LS, PVN and MeAD. Before implantation, the EAr was dipped in 1,1’-Dioctadecyl-3,3,3’,3’-tetramethylindocarbocyanine perchlorate (42364, Sigma-Aldrich) to visualize electrode locations post-mortem. The reference and ground wires were inserted into their respective burr holes and the EAr was lowered onto the surface of the exposed brain using a motorized manipulator (MP200; Sutter Instruments). The dorsoventral coordinates were estimated using the depth of the electrode targeting the PVN (AP = -1 mm, ML= -0.3 mm), which was slowly lowered to -4.7mm. Following its insertion at the desired location, the EAr and exposed skull with screws were secured with dental cement (Enamel plus, Micerium).

#### Neuropixels 1.0 probe implantation

For chronic electrophysiology experiments, animals were implanted with Neuropixels 1.0 probes (NP 1.0, Cat # PRB_1_4_0480_1_C, IMEC, Belgium). The probe was assembled in a custom-made 3D-printed probe holder, and the reference and ground wires were soldered to flex cable (silver wire, 127 µm, 300-500 Ω; A-M Systems, Carlsborg, WA) before implantation. Before surgery, the probe shank was soaked in a fluorescent Dil (42364, Sigma-Aldrich) for 10 min. Adult ICR male mice (32-38 g) were anesthetized under isoflurane (induction 3%, 0.5%-0.8% maintenance in 200 mL/min air; SomnoSuite) and placed over a custom-made heating pad (37°C) under a stereotaxic device (Kopf Instruments, Tujunga, CA). Two watch screws (0-80, 1/16”, M1.4) were inserted into the temporal bone to support the NP1 holder with dental cement. For implantation, the probe-containing holder was attached to a stereotaxic arm and centered at ML -0.4 mm, AP -1.9 mm, relative to bregma, after drilling the skull at this site to expose the dura. The DV coordinates at bregma were kept at 0.5 mm above the lambda to target the regions at an angle. The reference and ground wires were inserted into respective burr holes drilled into the right hemisphere. Then, the probe was lowered to DV -4.6 mm, relative to bregma, at a rate of 0.01 mm/s. A mix of petroleum oil and bone wax jelly gently filled the craniotomy, avoiding the shank to keep the tissue moist for long chronic recordings. Dental cement was applied to the exposed skull and base of the NP1 holder, along with screws to keep the probes firmly attached to the mouse head.

#### Recovery

After the surgery, antisedan (0.1ml/10g bodyweight) was given sub-cutaneously to wake the mice from the anesthesia. The mice were injected with meloxicam (5%, 0.01 mg/g) and baytril (5%, 0.03 ml/10g) to relieve pain and prevent infections for three days following surgery. Behavioral testing with optogenetic stimulation or fiber photometry was conducted three weeks after surgery to allow optimal viral expression. For electrophysiology recordings, animals were tested three days after surgery. All subject animals were singly housed following surgery so as to not disturb their implants.

### *In vivo* fiber photometry recordings

#### Test

All fiber photometry recordings were performed using the apparatus described above. The optic fiber of the implanted mouse was connected to the optical patch cords via a sleeve connector (Doric) under mild isoflurane anesthesia. The wired animal was then allowed to habituate in the behavior setup for 15 min. The recording session started with an additional 5 min baseline (pre-encounter) period with empty chambers. Thereafter, the empty chambers were replaced by chambers containing the appropriate stimuli for a 5 min test. Following the test, the stimuli-containing chambers were again replaced with empty ones for an additional 5 min of recordings. Each subject (n=13) animal was tested 2-3 times, with at least 40 min between distinct sessions with the same subject.

#### Fiber photometry recording and synchronization

Calcium signals were recorded using the RZ10x system of Tucker Davis Technologies and an optical path by Doric, which includes 600 μm mono-fiber optic patch cords connected to a four ports Fluorescence MiniCube (FMC4_IE(400-410)_E(460-490)_F(500-550)_S) and a 200 μm optical patch cord with a fiber-optic rotary joint (FRJ_1×1_PT_200/230/LWMJ-0.57_1m_FCM_0.15m_FCM) connected to the recorded animal. A camera (Flea3 USB3) was placed on top of the arena and connected to the digital port of the RZ10× system and configured to deliver strobes for every frame acquired at a frame rate of 30 frames/sec. These were later used to synchronize the video frames with the calcium signal. TDT Synapse software was used for recording the GCamp6f signal channel (excitation 465 nm, modulated at 330 Hz), the isosbestic control channel (405 nm, modulated at 210 Hz) and the digital channel receiving the camera strobes. LED currents were adjusted to return the voltage to between 0-200 mV for each signal, were offset by 5 mA, and were demodulated using a 6 Hz low-pass frequency filter.

### *In vivo* optogenetic stimulation

#### Test

In both the SP and ESPs tasks, subject animals (n=6) were first connected to the stimulation apparatus and left for 20 min of habituation to an arena containing empty chambers. The interaction was recorded for 5 min, after replacement of the empty chambers with the appropriate stimuli. Each subject was tested four times in each test, using a different stimulation protocol: Unstimulated, stimulated at the beginning of a social/stressed investigation bouts, stimulated at the beginning of an object/neutral investigation bouts, or stimulated consistently throughout the 5 min duration of the test. The four types of experiments were randomly conducted with the same animals, each on a separate day.

#### Optogenetic stimulation

Subject mice were lightly anesthetized with isoflurane and connected to the laser (473 nm, model FTEC2471-M75YY0, Blue Sky Research) via a sleeve connector (Thorabs, ADAF1) for light delivery. Animals were given 20-30 min to recover from any residual effects of the brief isoflurane anesthesia before testing, until they showed full mobility. Light power was set at 5 mW, which is sufficient to yield 1 mW/mm^2^, the minimum amount of light reported to be capable of efficient ChR2 activation ^48^, and positioned no farther than 0.25 mm above the mPFC. This was expected to specifically stimulate the mPFC, as previous measurements with blue light stimulations in rodent brains showed that the blue light of the laser does not penetrate the tissue deeper than 500 μm ^49^. Each animal underwent four sessions of SP and four sessions of ESPs each on a separate day, as described above. In all cases, the optogenetic stimulation was a 1 s train of 15 ms pulses delivered at 10 Hz. The optical stimulation was manually delivered via a Master8 Channel Programmable Pulse Stimulator (A.M.P.I).

### *In vivo* electrophysiological recordings

#### Test

All experiments were conducted in the experimental arena, as described above for the behavioral experiments. Before experiments, the mice were briefly exposed to isoflurane, and the microelectrode array (EAr) or Neuropixels 1.0 probe was connected to the evaluation system. Each recording session was divided into two 5 min stages, namely, a baseline period during which time the subject was alone in the arena in the presence of two empty chambers and a test (encounter) period when the subject performed the SP, ESPi or ESPs tasks. Each subject was tested over three sessions of the ESPi task, each conducted on a separate day.

#### EAr recordings

Electrophysiological recordings were conducted as previously described ^29,50^ via the RHD2000 evaluation system, using an ultra-thin SPI interface cable and a RHD2132 amplifier board (Intan Technologies). A custom-made Omnetics-to-MillMax adaptor was used to connect the electrode and amplifier. Recorded signals (sampled at 20 kHz) were synchronized with the video recording by a start signal sent through a custom-made triggering device and TTL signals from the camera to the recording system.

#### Neuropixels recordings

Data from the NP 1.0 probe was acquired at a 30 kHz sampling rate using SpikeGLX (https://billkarsh.github.io/SpikeGLX/) through a Neuropixels 1.0 PXIe acquisition system installed within a PXIe-1071 Express Chassis instrument (National Instruments). The camera timestamps strobes were recorded through a PXIe-6341 card (installed within the Chassis; National instruments) at 30 kHz for further alignment of the video clip to the physiological recordings.

### *Post-mortem* histological analysis of viral injection, optic-fiber and electrode placement

Each implanted mouse were anesthetized with isoflurane (induction 3%, 0.5%-0.8% maintenance in 200 mL/min air), attached to the stereotaxic device and the optic fiber/ Ear/NP1 probe was gently extracted from its head. The mouse was then perfused with phosphate buffer saline (PBS) and fixed using 4% paraformaldehyde (PFA, Sigma) solution. The brains were harvested and placed in PFA (4%) for 48 h, followed by sectioning of 50 mm slices in the horizontal axis using a VT1200s Leica sliding vibratome. The slices were collected onto microscope slides and stained with DAPI. Images of subject brain slices were acquired for verification of the viral expression placement and optic fiber path within the mPFC using an epifluorescence microscope (Ti2 eclipse, Nikon) equipped with a blue filter for DAPI staining, FITC for detecting GCaMP6-expressing cells, and TRITC for detecting ChR2-mCherry-expressing cells. EAr marks were visualized (DiI coated, Red) against DAPI-stained sections, and the marks were used to locate the respective brain regions based on the mouse atlas ^51^. Neuropixels probe tracks were aligned to the Allen mouse brain atlas as explained below.

## Quantification and statistical analysis

### Tracking software and behavioral analyses

All recorded video clips were analyzed using TrackRodent (https://github.com/shainetser/TrackRodent) ^46^. We used the *BlackMouseBodyBased* algorithm for C57BL/6J (black mice) and the *WhiteMouseBodyBased* algorithm for the ICR and *Shank3B* mutant mice (white mice). Similarly, for analysis of sessions with tethered animals (in the fiber photometry and optogenetics experiments), we used the *BlackMouseWiredBodyBased* and *whiteMouseWiredBodyBased* algorithms for white and black mice, respectively. Behavioral analysis was conducted as previously described ^52^.

### Synchronization of physiological data to behavioral events

We used a custom-made code called DeepPhenotyping to visualize and synchronize the different signals and behavioral events, as previously described ^28,53^. For each signal (spikes, LFPs or calcium signal) and event type (“All bouts”, “short bouts – <6 sec”, “long bouts – >6 sec”, “End of all bouts” and “End of long bouts – >6 sec”), the software considered the relevant sample rate and aligned the signal to the beginning of the event.

### Calcium signal data analysis

Calcium signals were analyzed using DeepPhenotyping. We first fitted the 405 channel onto the 465 channel to de-trend signal bleaching and any movement artifacts, according to the manufacturer’s protocol (https://github.com/tjd2002/tjd-shared-code/blob/master/matlab/photometry/FP_normalize.m). Next, the de-trended signal was aligned to the video recording using the timestamps recorded by the digital port of the RZ10× system. We aligned the de-trended signal to each behavioral event (short/long bouts) and normalized it using the Z*-*score (0.5 s bins), with the pre-event period of -4 s to -1 s serving as a baseline.

### Analysis of LFP signals

All signals were analyzed using DeepPhenotyping, as previously described ^50^. First, the signals were down-sampled to 5 kHz and low-pass filtered up to 0.3 kHz using a Butterworth filter. The power over time for the different frequencies was constructed with the ‘‘spectrogram” function in MATLAB, using a 2-s long discrete prolate spheroidal sequence (DPSS) window with 50% overlap, at 0.5 Hz increments and 0.5 s time bins. The power for each frequency band (Theta: 4– 12 Hz and Gamma: 30–80 Hz) was averaged, and the delta in dB was calculated, relative to the mean power averaged across the entire 5 min of the pre-encounter period. Changes in theta (ΔθP) and gamma (ΔγP) power for each brain region were defined as the mean difference in power between the encounter and pre-encounter periods. Synchronization between the LFP signal and investigation bouts towards the stimuli was assessed by calculating the gamma power within a time window of 5 s before and 5 s after the beginning of each bout and normalizing it using Z-score (0.5 s bins) analysis, with the pre-event period -5 s to 0 s serving as a baseline.

### Post-process of Neuropixels 1.0 data

Spiking data of the behavior experiments acquired from SpikeGLX, were then sorted into spike clusters with Kilosort 2.5 (https://github.com/MouseLand/Kilosort). These sorted spikes were further manually curated to identify single units, separate them from multi-unit spikes and remove noise clusters using phy (https://github.com/cortex-lab/phy). Single units were manually curated according to the following criteria: Less than 0.1% of spikes violated the cell refractory period of 2 ms) and spike waveform was consistent with a single unit. Waveform and cross-correlograms of all nearby units were compared to verify that no two clusters corresponded to the same neuron. The recorded units were registered with respective brain regions in accordance with Allen’s brain mouse atlas using Universal probe finder ^54^. The original code was modified to extract the spikes data, and camera strobe timestamps were further analyzed by DeepPhenotyping (10.5281/zenodo.10708903).

### Analysis of single-unit responses to behavior events

To calculate the neuronal response to each behavioral event, the mean activity for each neuron across all similar events in the same session was quantified and Z-scored based on the mean and standard deviation for this neuron at various baseline periods, according to the desired analysis (for Figs. 8 and S6, -4 s to -1 s; for Figs. 9 and S7, -2 s to -1 s). For hierarchical clustering, each neuron’s Z-score response (begin and end of bouts of investigation of each stimulus) was horizontally concatenated and then clustered using agglomerative hierarchical clustering with Euclidean distance (100 au) as a threshold, which resulted in the separation of 6-7 functional groups per task, using a custom-written Python script (10.5281/zenodo.10708903). Unbalanced data due to missing investigation of one of the stimuli were filled with zeros to keep the matrix structure appropriate for clustering. Furthermore, the percentage of neurons that were excited (> 1.9 Z-score) or inhibited (-<1.9) in specific events for each task was estimated from the mean firing rate (Z-score) from time 0 to 2 seconds after the event began or ended.

### Logistic regression classifier

To test whether PFC or ACC neural activity within each group was correlated with the dynamics of an investigation bout directed towards a specific event (begin or end of bouts) and stimulus type, we employed logistic regression classifiers with L2 regularization ^55,56^. The neural activity within each time bin for each area and clustered neuronal group (comprising at least 20 neurons) was divided into five non-overlapping parts to allows the use of Kfolds cross-validation. The classifier was trained and tested on the training folds to predict one of the two stimuli and evaluated on the held-out data. This step was iterated 100 times to normalize the random initiation of Kfolds, and the prediction accuracy was averaged over these iterations for each time-bin. The same process was repeated with shuffled data to evaluate whether these models were not over-fitting, and the results were compared.

### Statistical analysis

All results and parameters of the statistical analyses are detailed in Table S1 according to the various figures. The sample size for all behavioral experiments was based on previously published power calculations ^46^. All statistical tests were performed using SPSS v26.0 (IBM) or GraphPad Prism (v10.1.2). The Kolmogorov–Smirnov and Shapiro-Wilk tests were used for verifying the normal distribution of the dependent variables. A two-tailed paired t-test was used to compare different conditions or stimuli for the same group, and a two-tailed independent t-test was used to compare a single variable between distinct groups. When the normal distribution assumption was violated, the non-parametric Wilcoxon test was used to compare different conditions or stimuli for the same group, and a Mann-Whitney was used to compare different conditions or stimuli for different groups. For comparison between multiple groups and parameters, a mixed model (MM) analysis of variance (ANOVA) was applied to the data. This model contained one random effect (ID), one within effect, one between effect, and the interaction between them. For comparison within a group using multiple variables, a two-way repeated-measures (RM) ANOVA model was applied to the data. This model contained one random effect (ID), two within effects, one between effect, and the interactions between them. All ANOVA tests were followed, if main effects or interaction found, by a *post-hoc* Student’s t-test with Holm-Sidak correction or FDR corrections, with Q < 0.001. When the sphericity assumption needed for ANOVA was violated, the Greenhouse-Geisser correction was used. Significance was set at 0.05 and was adjusted using Holm-Sidak or FDR correction when multiple comparisons were used.

## Supplementary figures

**Figure S1.**
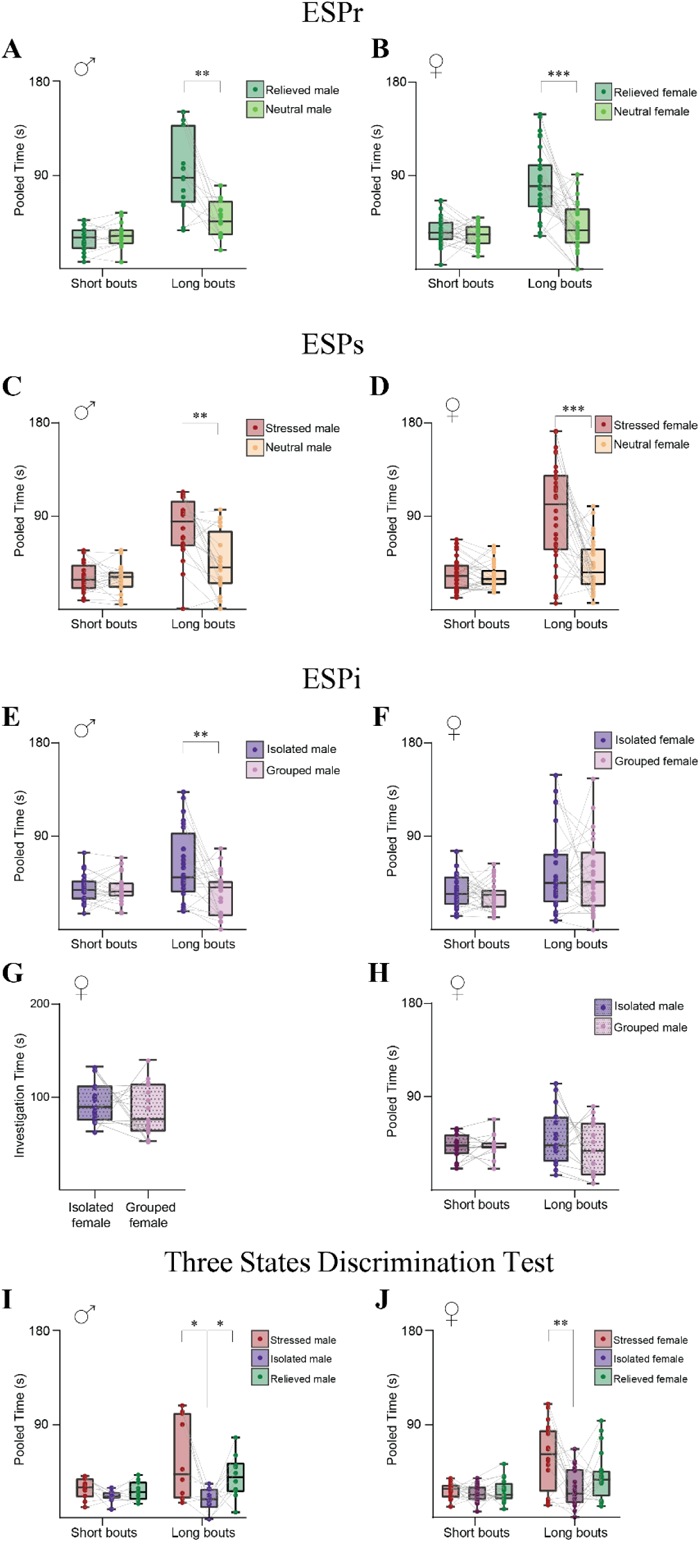
Short and long investigation bouts during the various behavioral tasks. **A.** Mean time of short (left) and long (right) events dedicated by male subjects for investigation of relieved and neutral (see legend) stimulus animals during the ESPr task. A significant effect was found for the stimulus X bout duration interaction (*p*<0.01) by a two-way RM ANOVA. **B.** As in **A**, for female subjects. A significant effect was found for the stimulus X bout duration interaction (*p*<0.01) by a two-way RM ANOVA. **C-D.** As in **A-B**, for the ESPs task. *Males*: A significant effect was found for the stimulus X bout duration interaction (*p*<0.01) by a two-way RM ANOVA; *Females*: A significant effect was found for the stimulus X bout duration interaction (*p*<0.01) by a two-way RM ANOVA. **E-F.** As in **A-B**, for the ESPi task. *Males*: Significant effects were found in stimulus (*p*<0.01) and bout duration (p<0.01) by two-way RM ANOVA were found for; *Females*: A significant effect was found for the stimulus X bout duration interaction (*p*<0.05) by a two-way RM ANOVA. **G.** Mean time dedicated by female subjects for investigating isolated and group-housed male (see legend) stimulus animals during the ESPi task. **H.** As in **G**, divided into long and short bouts. **I-J.** As in **A-B**, for the three-state discrimination task. *Male:* Long Bouts – A significant main effect was found for stimulus (*p*<0.05) by one-way RM ANOVA; *Females*: Long Bouts-A significant main effect was found for stimulus (*p*<0.01) by one-way RM ANOVA. *p<0.05, \*\**p*<0.01, \**p*<0.05, *post-hoc* paired LSD comparisons following detection of a main effect by ANOVA.

**Figure S2.**
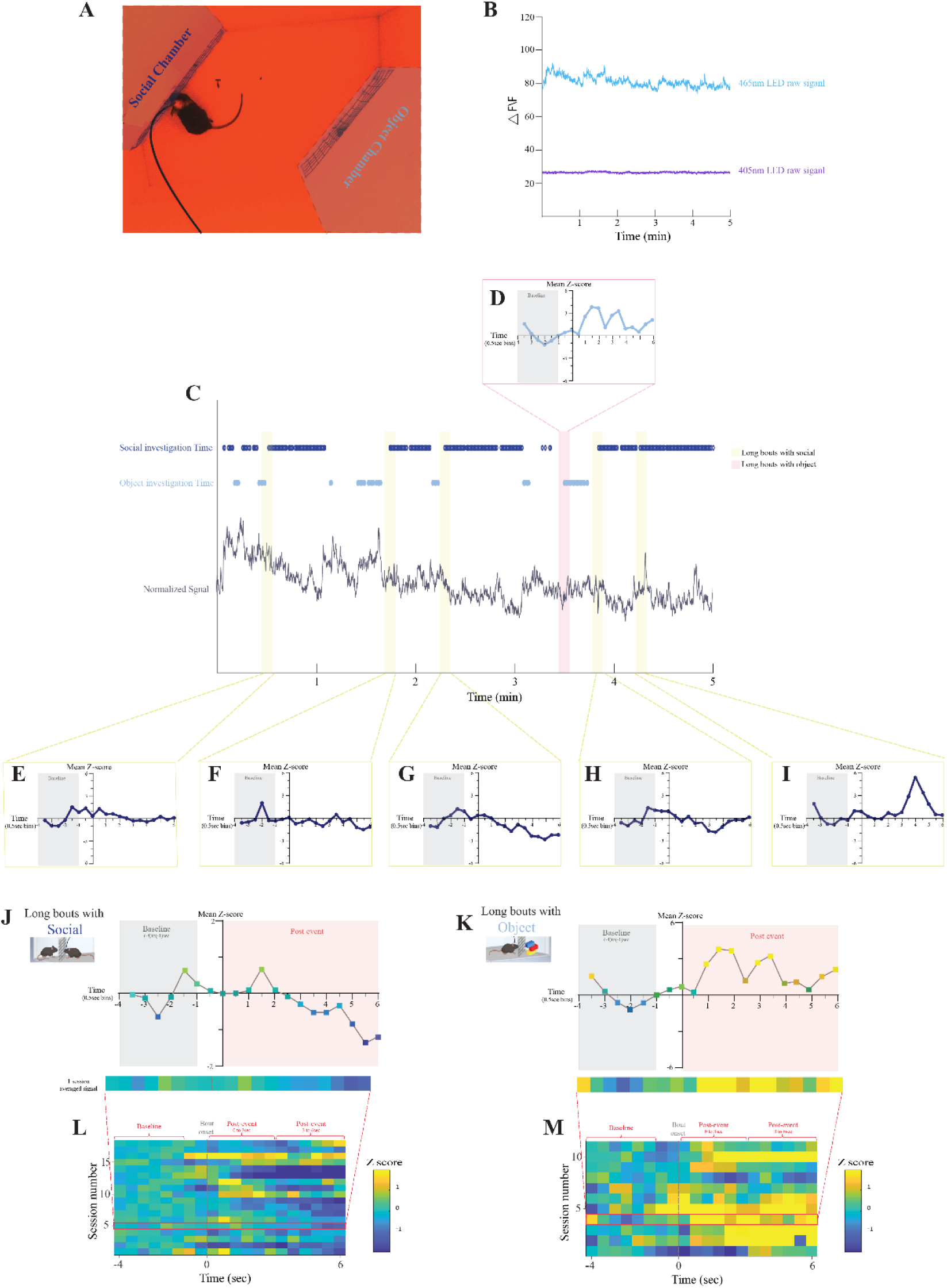
Analysis pipeline of fiber-photometry calcium signals. **A.** A picture taken from above the subject mouse in the arena during a fiber photometry recording session. **B.** Example raw traces of the signals recorded via the 465 nm (light blue) and 405 nm (purple) channels, during a single 5 min test stage of the SP task. **C.** The normalized signal after fitting and subtraction of the 405 nm channel signal from the 465 nm signal, shown together with the social (blue) and object (light blue) investigation bouts (see dots above). Pink and yellow bars mark long investigation bouts towards the object (pink) and social (yellow) stimuli taken from the session for Z-score analysis. **D.** A Z-scored trace of the calcium signal shown in **C**, recorded four seconds before and six seconds after the beginning of a long investigation bout towards the object. **E-I.** As in **D** for several long investigation bouts towards the social stimulus in the same session. **J.** Mean Z-scored signals for the long social investigation bouts, generated by averaging all the traces shown in **E-I**. A heat-map representing the trace is shown below **K.** As in **J**, for the object stimulus. **L.** Combined heat-map for long social investigation bouts across all SP sessions. **M.** As in **L**, for object investigation bouts.

**Figure S3.**
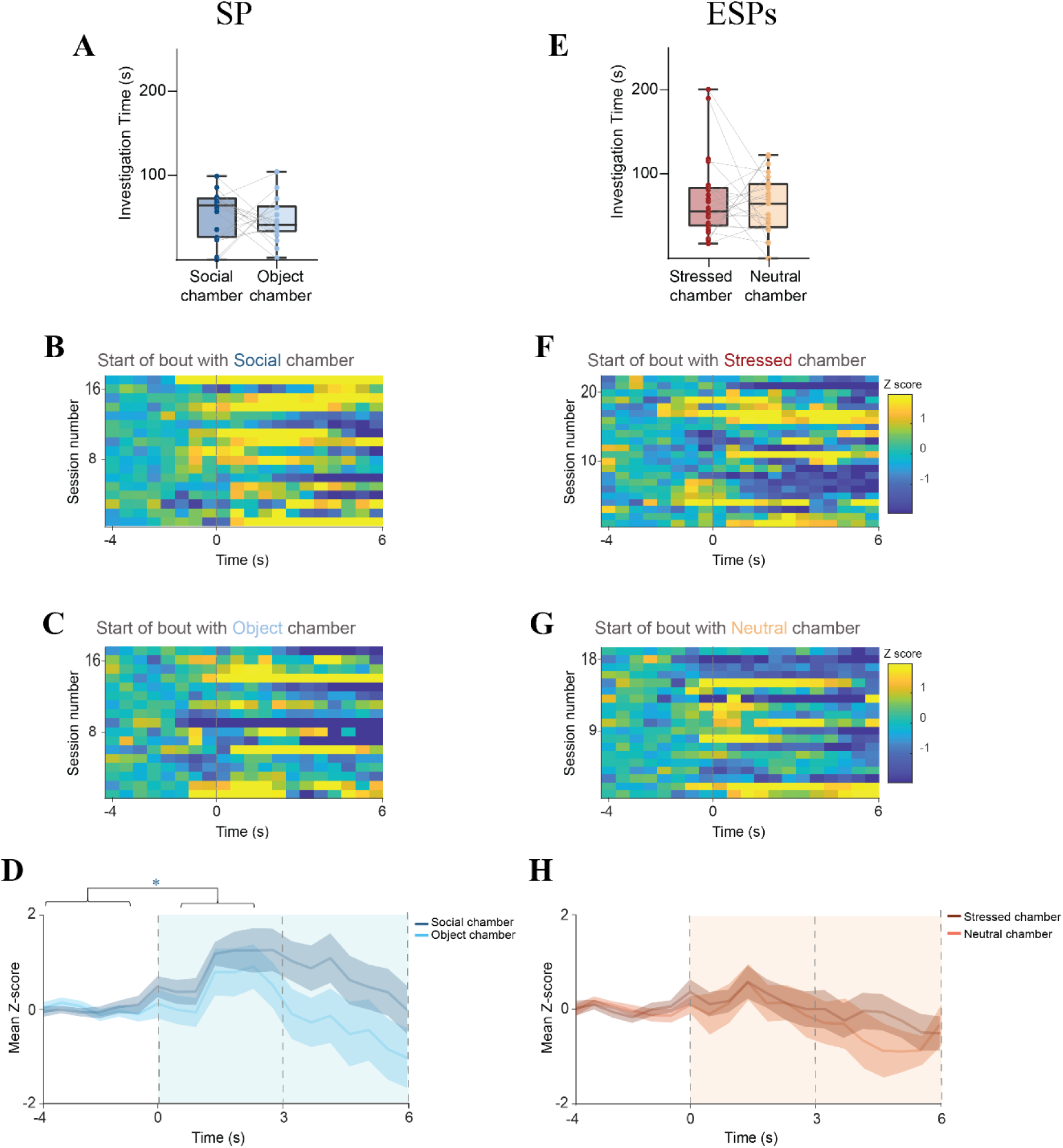
No chamber-associated differences in calcium signals were recorded during investigation of empty chambers. **A.** Mean time dedicated by recorded male subjects for investigating empty chambers to be later used to house the social (blue) and object (light blue) stimuli during the 5-min pre-encounter stage of the SP task. No significant difference was found between times in the two chambers in a paired-samples t-test. **B.** Heat-map (see color code to the right of **F**) of the Z-scored calcium signals at the beginning (first six seconds) of bout, averaged across all investigation bouts towards the empty chamber to be later used for housing the social (blue) stimulus during the 5 min pre-encounter stage of the SP task (each line corresponds to a single session), using 0.5-s bins. Time ‘0’ represent the beginning of the bout. **C.** As in **B**, for the empty chamber be be later used for housing the object stimulus in the same sessions. **D.** Super-imposed traces of the mean (±SEM) Z-scored calcium signals shown in **A-B**, averaged across all sessions. Dashed lines represent three distinct time windows of the event, which were used for the statistical analysis. A significant effect was found for the time interaction (*p*<0.05) by two-way MM ANOVA. **E-H.** As in **A-D**, for the ESPs task \**p*<0.05, \*\**p*<0.01, \*\*\**p*<0.001, *post-hoc* paired (within stimulus differences in time) t-test with Holm-Sidak correction for multiple comparisons following the detection of main effects by ANOVA.

**Figure S4.**
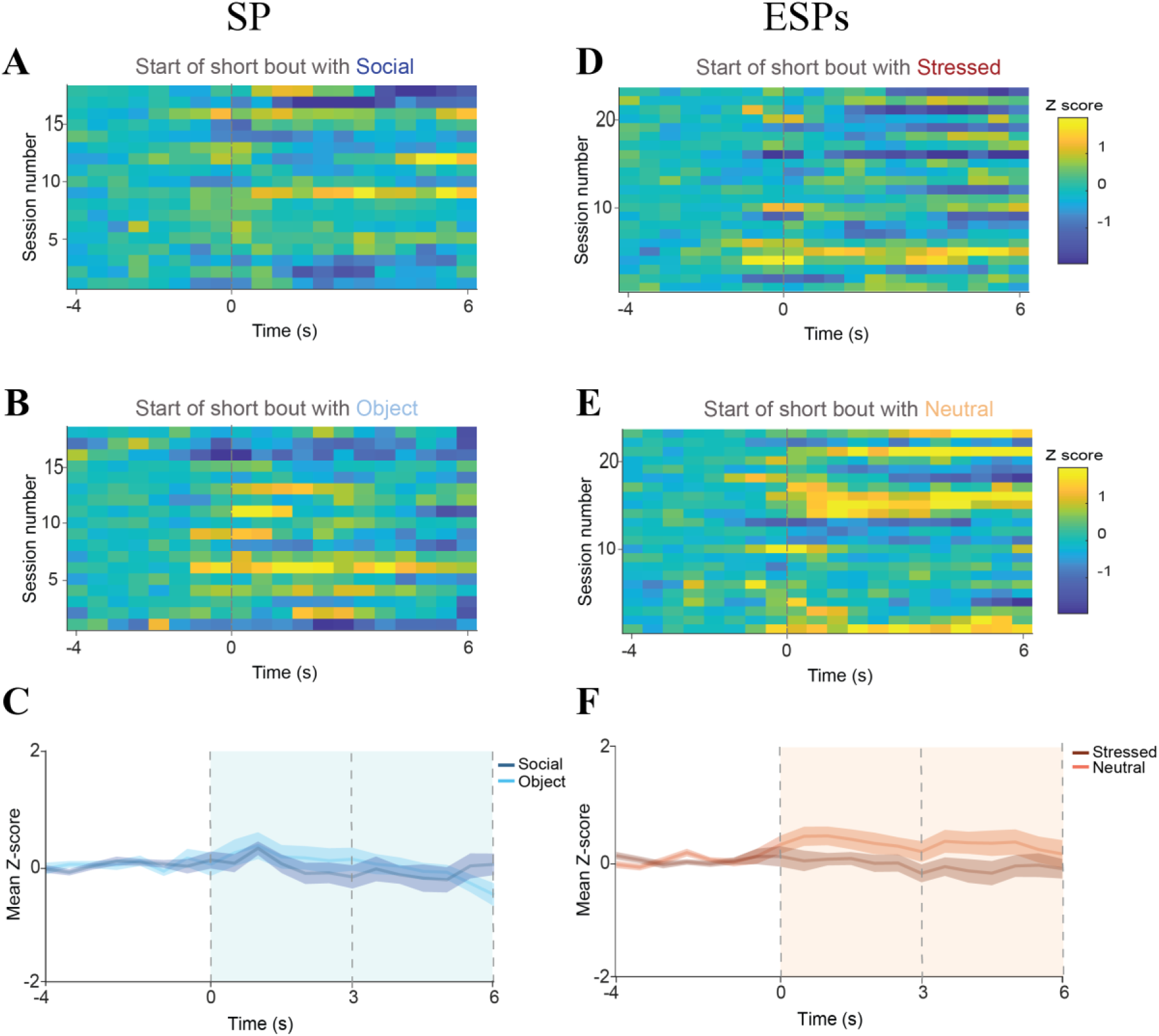
Calcium signals recorded from *CamK2a*-positive mPFC neurons in short investigation bouts during the SP and ESPs tasks. **A.** Heat-map (see color code to the right of **B**) of the Z-scored calcium signals at the beginning (first six seconds) of bout, averaged across all short (<6 s) investigation bouts towards the stimulus animals, shown for all sessions of the SP task (each line corresponds to a single session), using 0.5-s bins. Time ‘0’ represent the beginning of the bout. **B.** As in **A**, for the object stimulus in the same experiments. **C.** Super-imposed traces of the mean (±SEM) Z-scored calcium signals shown in **A-B**, averaged across all sessions. Dashed lines represent three distinct time windows of the event, which were used for the statistical analysis. No significant effects were found between the stimuli and time windows by two-way MM ANOVA. **E-H.** As in **A-D**, for the ESPs task. No significant effects were found by two-way MM ANOVA.

**Figure S5.**
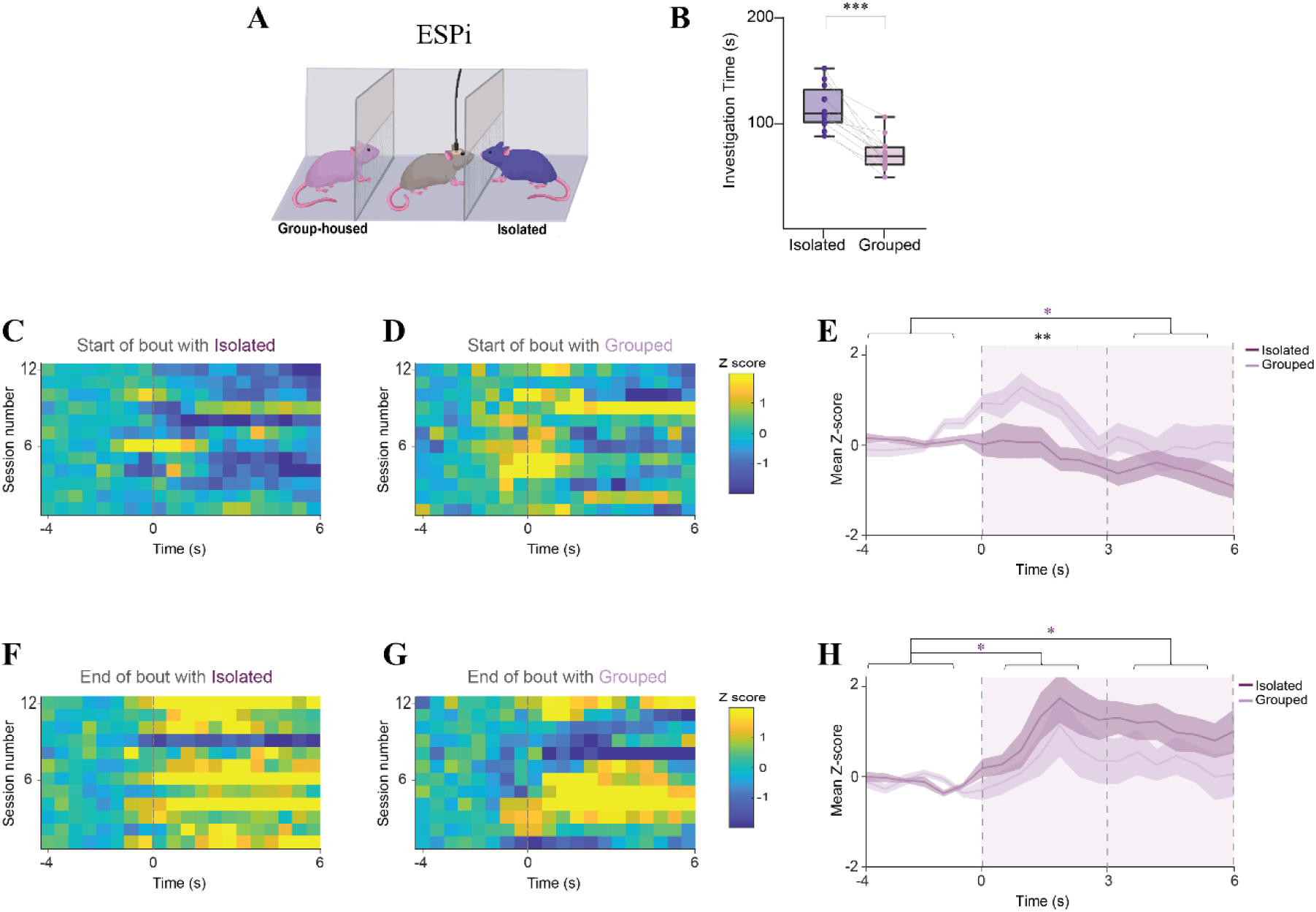
Calcium signals recorded from *CamK2a*-positive mPFC neurons in long investigation bouts during the ESPi task. **A.** Schematic representation of the ESPi task. **B.** Mean investigation time towards the isolated (purple) or group-housed (light purple) stimulus animals during the ESPi tasks (n=12 sessions from 8 subjects) conducted with fiber photometry. *\*\*\*p<*0.001, paired samples t-test. **C.** Heat-map (see color code to the right of **D**) of the Z-scored calcium signals at the beginning (first six seconds) of bout, averaged across all long investigation bouts towards the stimulus animals, shown for all sessions of the ESPi task (n=12 sessions, each line corresponds to a single session), using 0.5-s bins. Time ‘0’ represent the beginning of the bout. **D.** As in **D**, for the group-housed stimulus animal in the same experiments (n=12 sessions). **E.** Super-imposed traces of the mean (±SEM) Z-scored calcium signals shown in **C-D**, averaged across all ESPi sessions. Dashed lines represent three distinct time windows of the event, which were used for the statistical analysis. Asterisks, representing a statistical comparison between the stimuli and across time windows, are color-coded according to the type of comparison. Purple and light purple asterisks represent a significant difference for social and object investigation, respectively, during the specific time window, as compared to baseline, while black asterisks represent a difference between the two stimuli during the same time window. Significant effects were found for time (*p<*0.05) and stimulus (*p*<0.05) by two-way MM ANOVA. **F-H.** As in **C-E**, for the end of long investigation bouts. A significant effects was found for time (*p*<0.01) by two-way MM ANOVA. \**p*<0.05, \*\**p*<0.01, *post-hoc* paired (within stimulus differences in time) or independent (differences between stimuli within time windows) t-test with Holm-Sidak correction for multiple comparisons following detection of main effects by ANOVA.

**Figure S6.**
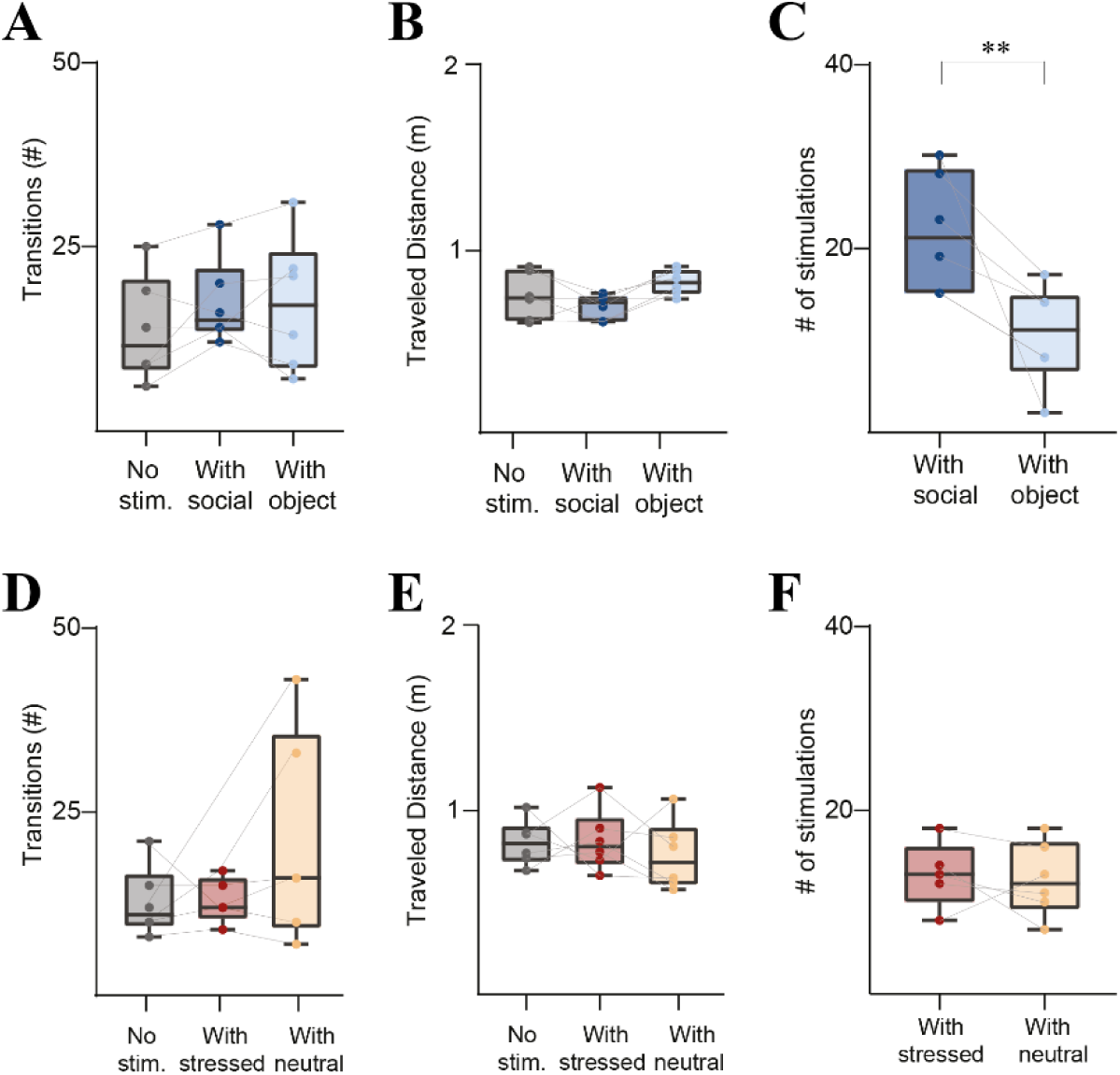
Behavioral and stimulation variables for the various stimulation protocols applied during the SP and ESPs tasks. **A.** Mean number of transitions of the two stimuli, made by subjects during the SP task with either no optogenetic stimulation (grey), or 1-s stimulation at the beginning of investigation bouts towards the social (blue) or object (light blue) stimulus. **B.** As in **A**, for the distance travelled by the subjects. **C.** Mean number of optogenetic stimuli applied at the beginning of investigation bouts towards the social (blue) or object (light blue) stimulus. Note that there were more stimuli with the social stimulus, due to significantly higher number of investigation bouts towards social stimuli, as compared to object stimuli. \*\**p*<0.01, independent samples t-test. **D-F.** As in **A-C**, for the ESPs. Note the lack of significant difference in the number of stimuli applied during investigation of stressed (left) and neutral (right) stimulus animals.

**Figure S7.**
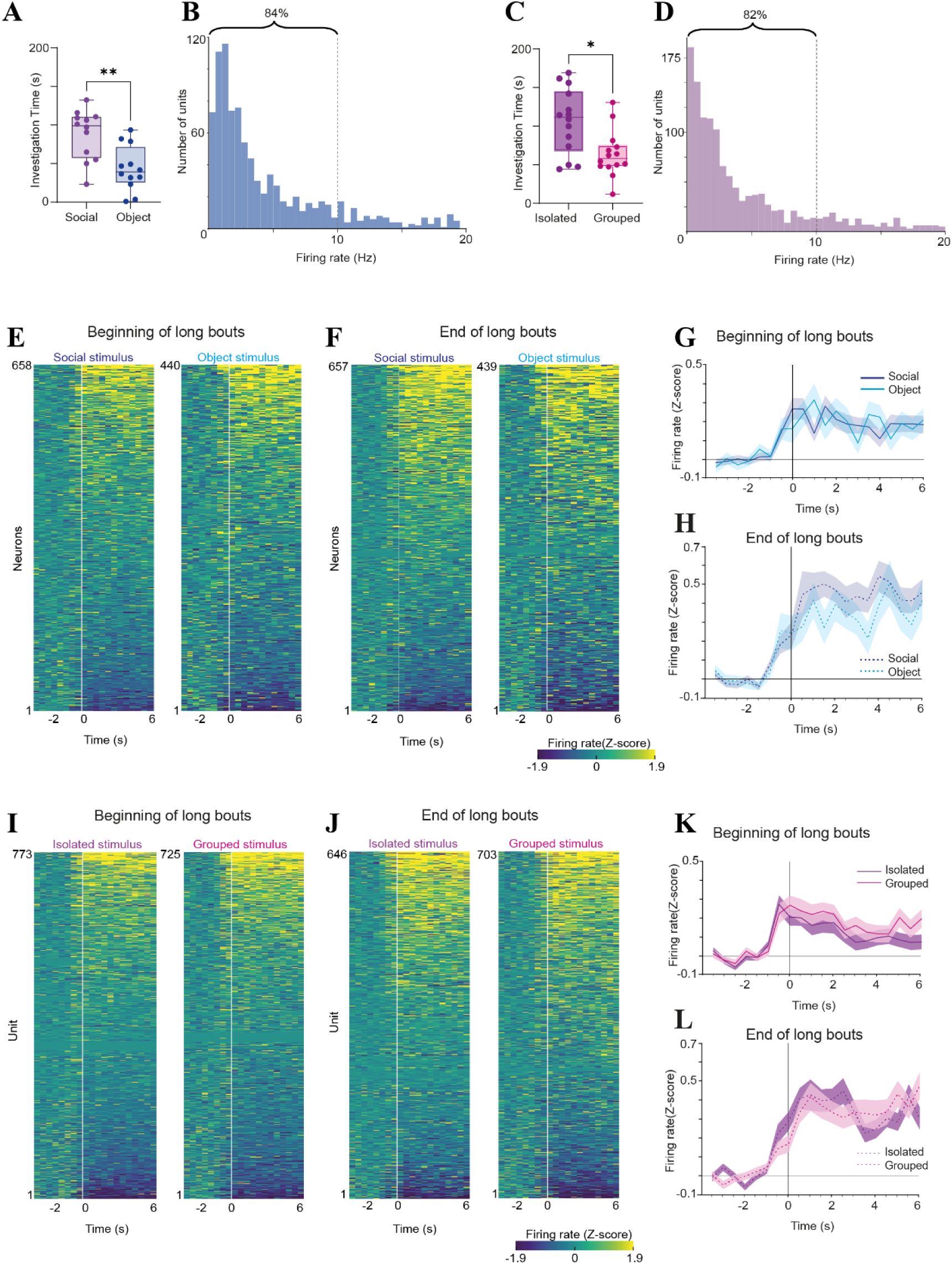
Responses of prefrontal cortical neurons to the two stimuli during the SP and ESPi tasks. **A.** Median investigation time towards the social (left) and object (right) stimuli during the SP session conducted by Neuropixels 1.0-implanted subject mice. Wilcoxon matched pairs signed rank test, n = 12 sessions, W = -68, \**p* =0.0049. **B.** Distribution of the of the number of recorded units according to their firing frequency during the baseline (pre-encounter) period (0.5 Hz bins) of SP task sessions. Note that 84% of the units fired below 10 Hz. **C.** As in **A**, for the stressed (left) and neutral (right) stimulus animals during the ESPs task, n = 13 sessions, W = -59, \**p* =0.0398. **D.** As in **B**, for ESPi sessions. **E.** Heat-maps of the mean Z-scored firing rate recorded at the beginning of long investigation bouts towards the social (left) and object (right) stimulus animals, for all units that fired at least one spike during this period of the SP task (hence the distinct number of units for each stimulus animal). The various units are arranged according to their responses from strongest excitation (above) to strongest inhibition (below), separately for each stimulus animal. **F.** As in **C**, for end of bout. **G.** Super-imposed traces of the mean Z-scored firing rate of all neurons shown in **C**. No statistically significant difference was observed (multiple unpaired t-test for each 0.5 s bin, followed by BHFDR correction). **H.** As in **G**, for all neurons shown in **D**. **I-L.** As in **E-H**, for the ESPi task.

**Figure S8.**
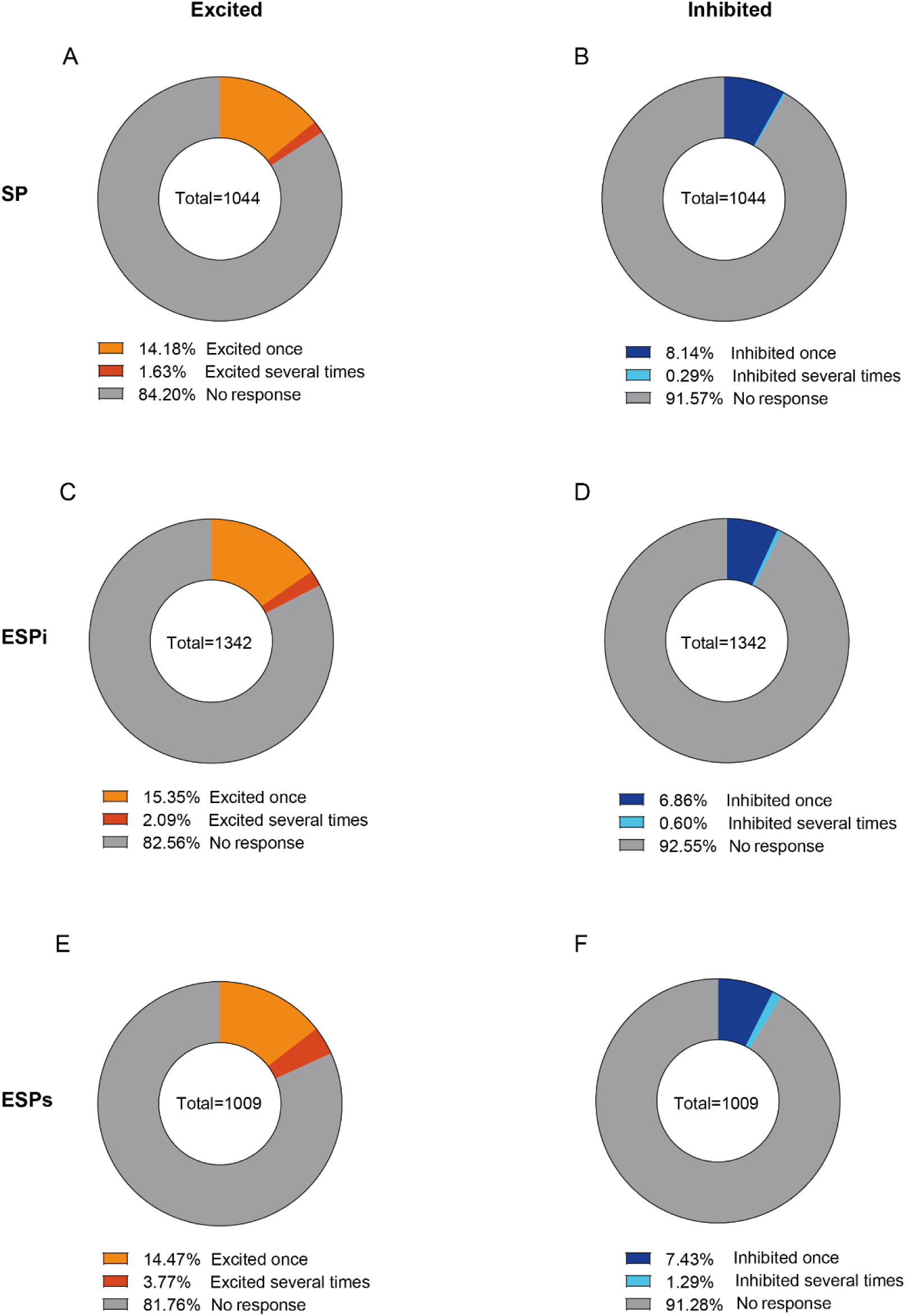
Most responsive neurons are event-specific. **A.** Circular distribution of all neurons recorded during SP sessions divided into three groups: Grey-neurons which did not pass the criteria for an excitatory response (mean Z-score of 1.9 across the first two seconds of either the beginning or end of all bouts of investigation of either the social or object stimulus), orange - neurons which responded with excitation during only one event, and red - neurons which responded with excitatory response during more than one event. **B.** As in **A**, for neuron showing inhibitory responses (mean Z-score of -1.9) during the SP task. **C-D.** As in **A-B**, for the ESPi task. **E-F.** As in **A-B**, for the ESPs task.

**Figure S9.**
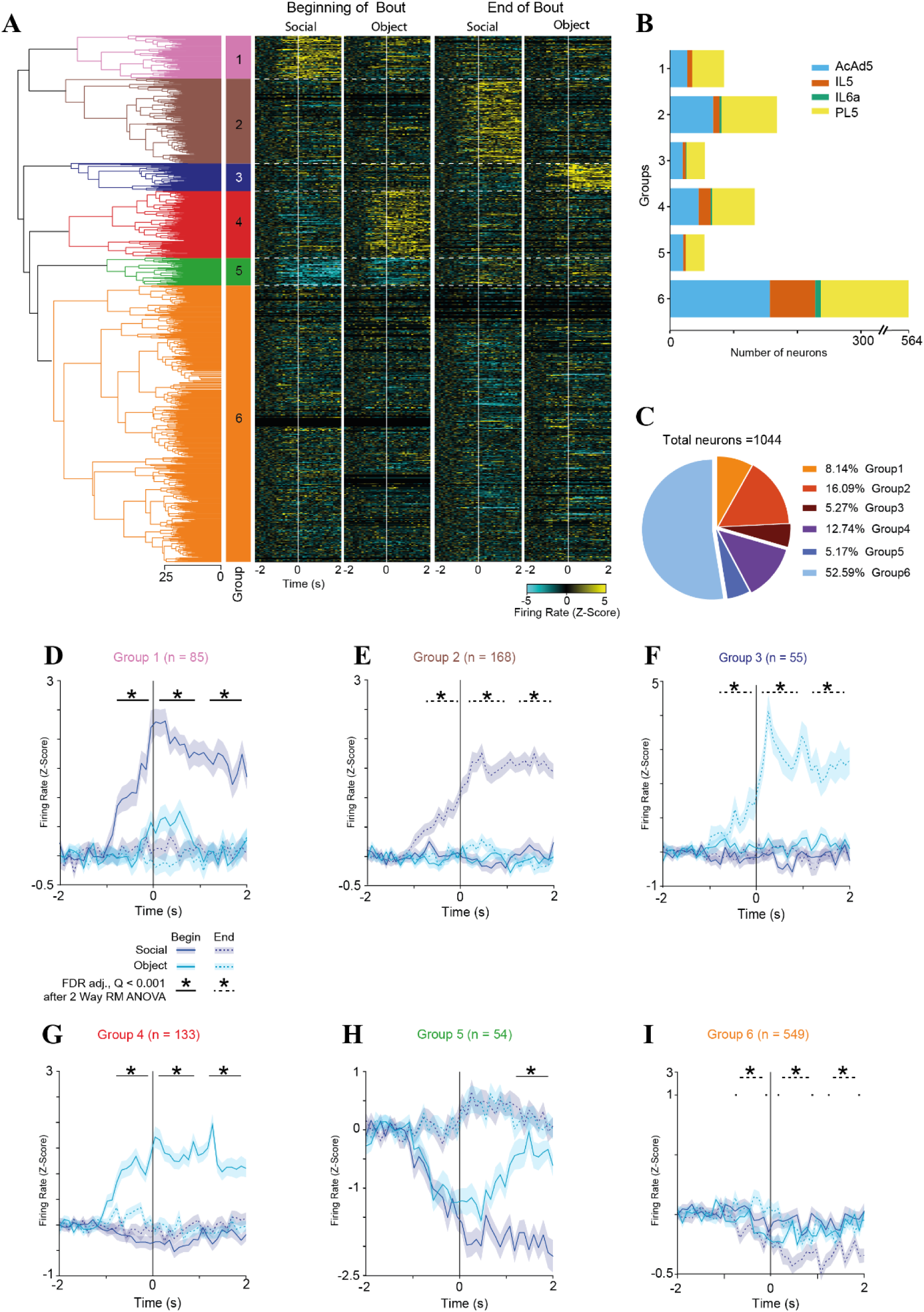
Distinct groups of prefrontal neurons differentially respond to the distinct stimuli during the SP task. **A.** A dendrogram of a hierarchical clustering of all cells that fired during either the beginning or end of investigation bouts towards any of the stimuli during the SP task, according to their Z-scored response. A heat-map showing the mean firing response of each cell during all four event types (denoted above) is displayed to the right of the dendrogram. **B.** A color-coded distribution of the units comprising each group to four recorded prefrontal regions, as detailed in the legend to the right: AcdAd5 – layer 5 of the dorsal anterior cingulate cortex, IL5 – layer 5 of the infralimbic cortex, IL6a - layer 6a of the infralimbic cortex, and PL5 – layer 5 of the prelimbic cortex. **C.** A color-coded pie chart showing the fraction (in %) of each group among all recorded units. **D.** Super-imposed traces of the mean Z-scored firing rate of the units included in Group 1 at the beginning (solid lines) and end (dashes lines) of bouts of investigation of social (purple) and object (light blue) stimuli. \**p*<0.001 in FDR-adjusted *post-hoc* t-test between the responses (averaged at 1-s bins) at the beginning of investigation bouts for the two stimuli, following a two-way RM ANOVA between time bins and stimulus. Similarly, for end of investigation bouts, \**p*<0.001 FDR adjusted *post-hoc* test following main effects in a comparison between time bins and stimulus. Note that the differential response started as early as one second before investigation. **E-I.** As in **D**, for Groups 2, 3, 4, 5 and 6, respectively.

**Figure S10.**
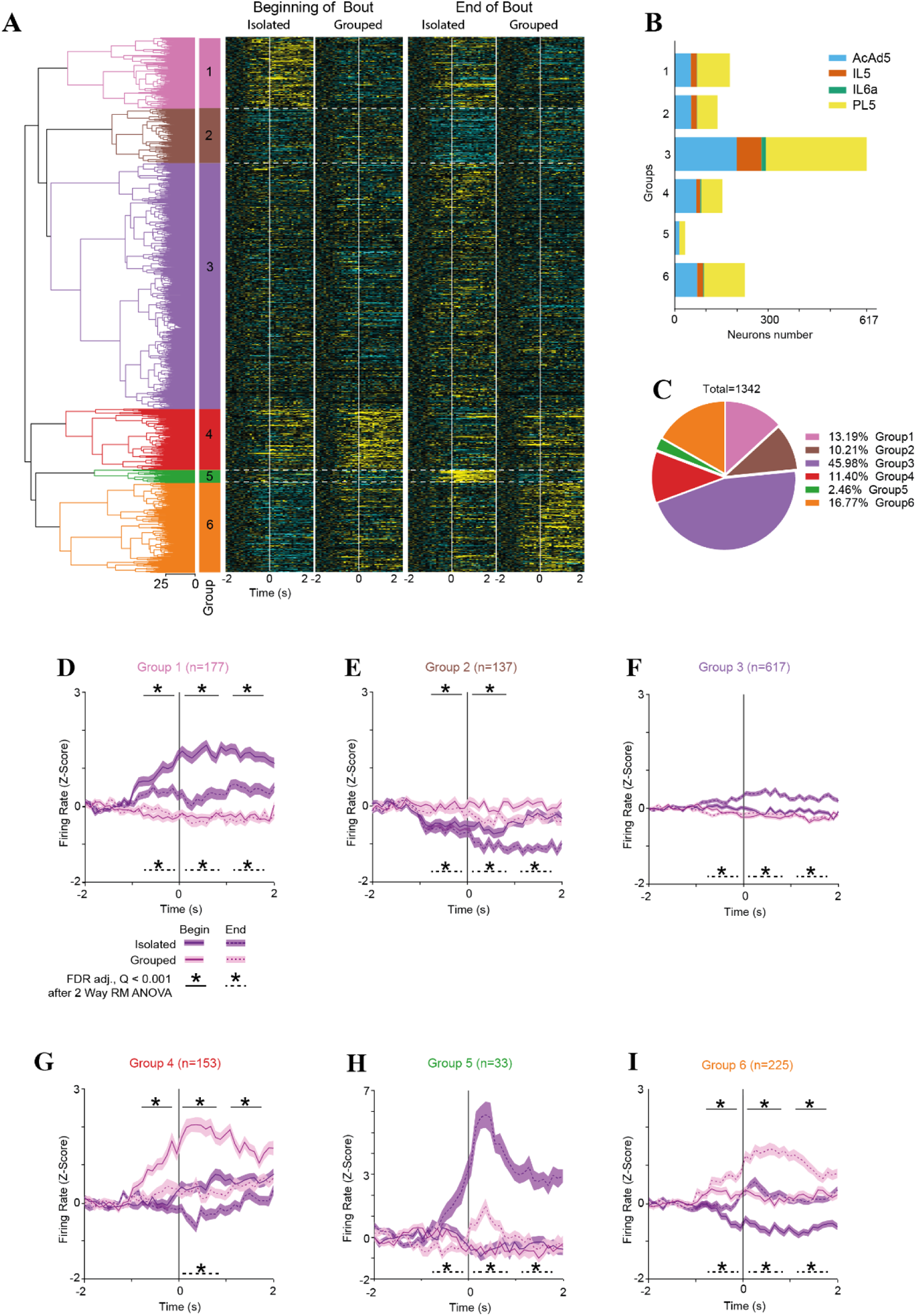
Distinct groups of prefrontal neurons differentially respond to the distinct stimuli during the ESPi task. **A.** A dendrogram of a hierarchical clustering of all cells that fired during either beginning or end of investigation bouts towards any of the stimuli during the ESPi task, according to their Z-scored response. A heat-map showing the mean firing response of each cell during all four event types (denoted above) is displayed to the right of the dendrogram. **B.** A color-coded distribution of the units comprising each group to four recorded prefrontal regions, as detailed in the legend to the right: AcdAd5 – layer 5 of the dorsal anterior cingulate cortex, IL5 – layer 5 of the infralimbic cortex, IL6a -layer 6a of the infralimbic cortex, and PL5 – layer 5 of the prelimbic cortex. **C.** A color-coded pie chart showing the fraction (in %) of each group among all recorded units. **D.** Super-imposed traces of the mean Z-scored firing rate of the units included in Group 1 at the beginning (solid lines) and end (dashes lines) of bouts of investigation of isolated (purple) and group-housed (pink) stimuli. \**p*<0.001 in FDR-adjusted *post-hoc* t-test between the responses (averaged at 1-s bins) at the beginning of investigation bouts for the two stimuli, following a two-way RM ANOVA between time bins and stimulus. Similarly, for end of investigation bouts, \**p*<0.001 FDR adjusted *post-hoc* test following main effects in a comparison between time bins and stimulus. Note that the differential response started as early as one second before investigation. **E-I.** As in **D**, for Groups 2, 3, 4, 5 and 6, respectively.

**Figure S11.**
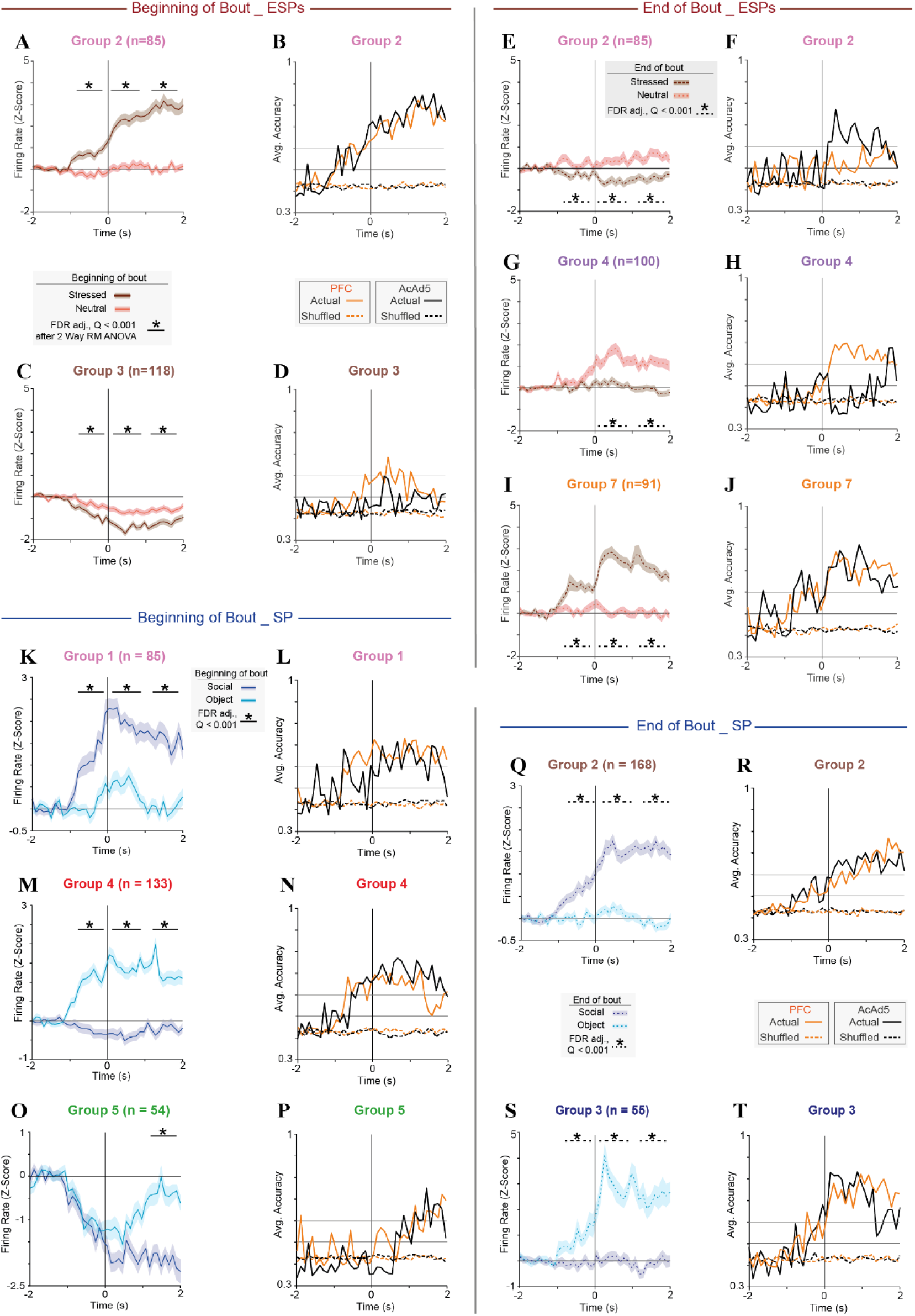
Activity of prefrontal neurons accurately discriminates between stimuli as early as one second before the beginning or end of investigation bouts. **A.** Super-imposed traces of the mean (±SEM) Z-scored firing rate of neuronal group 2 during the ESPs task, at the beginning of investigation bouts towards either the stressed (brown) or neutral (red) stimulus animal. Each asterisk represents 1 sec of significant difference between the stimuli. **B.** Traces representing the accuracy of a logistic model predicting the identity of the investigated stimulus according to the firing rate shown in **A**, analyzed separately for mPFC (orange trace) and ACC (black trace) neurons. The results of the same model using shuffled data are shown as dashed lines, using the same colors. Note that the predictive accuracy started to rise about 1 s before the beginning of the bout, for both mPFC and ACC neurons. **C-D.** As in **A-B**, for neuronal group 3. **E-F.** As in **A-B**, for the end of investigation bouts. Note that only ACC, but not mPFC neurons showed high (≥0.7) predictive accuracy. **G.H.** As in **E-F**, for neuronal group 4. Note that in the group, only mPFC, but not ACC neurons showed high predictive accuracy. **I-L.** AS in **E-F**, for neuronal group 7. Note that the predictive accuracy started to rise about one second before the end of the bout, for both mPFC and ACC neurons. **K-L.** As in **A-B**, for neuronal group 1 during the SP task, at the beginning of investigation bouts towards either the social (purple) or object (light blue) stimulus animal. **M-N.** As in **K-L**, for neuronal group 4. **O-P.** As in **K-L**, for neuronal group 5. Note that despite the significant inhibition in firing rate displayed as early as 1 s before the beginning of bout, the predictive value for both mPFC and ACC neurons remained low until 1 s after the beginning of bout, due to the similar reaction of the neurons to both stimuli to that point. **Q-R.** As in **K-L**, for neuronal group 2 of the SP task. **S-T.** As in **Q-R**, for neuronal group 3.

